# Chromatin dynamics controls epigenetic domain formation

**DOI:** 10.1101/2021.01.18.427115

**Authors:** Marina Katava, Guang Shi, D. Thirumalai

## Abstract

In multicellular organisms, nucleosomes carry epigenetic information that defines distinct patterns of gene expression, which are inherited over multiple generations. The enhanced capacity for information storage arises by nucleosome modifications, triggered by enzymes. Modified nucleosomes can transfer the mark to others that are in proximity by a positive feedback (modification begets modification) mechanism. We created a polymer model in which each bead, representing a nucleosome, stochastically switches between unmodified (*U*) and modified (*M*) states depending on the states of the neighbors. Spreading is initiated at a specific nucleation site (NS) that is permanently in the *M* state. Modifications spread among the non-nucleation loci probabilistically. If the spreading rate is higher than the chromatin relaxation rate, domains containing the modified nucleosomes form without bound. In the opposite biologically relevant limit, finite-sized domains form, driven by contacts between nucleosomes through a three-dimensional looping mechanism, with chromatin remaining in an expanded state, as is appropriate for fission yeast. Surprisingly, finite bounded domains arise *without* the need for any boundary elements as long as the spreading is slow. Maintenance of spatially and temporally stable domains require the presence of the NS whose removal eliminates finite-sized modified domains. By varying the solvent quality we show that finite heterochromatin domains form as long as initially chromatin is not fully condensed. The predictions compare well with experimental data for H3K9me3 spreading in Mouse Embryonic Stem cell.

## I. INTRODUCTION

The inheritance of distinct phenotypes that are not encoded in the DNA sequence has been demonstrated in multicellular organisms. The resulting distinct morphological characteristics are maintained over multiple cellular divisions [1, 2]. Alterations in the genetic material, resulting in distinct gene expression and subsequent phenotype variations, without any change to the underlying DNA sequence, are referred to as epigenetic modifications, which are carried over multiple cell divisions. As a consequence, some aspects of cellular memory are often associated with the term epigenetics [3–5]. The study of the establishment and inheritance of genetic patterns constitutes a burgeoning field, especially because epigenetic misregulation is implicated in aging and cancer. In eukaryotes, DNA condenses to form chromatin by wrapping around histone proteins. The physicochemical mechanisms governing genetic activity constitute a myriad of strategies that change the structure and organization of chromatin. These include chemical tagging of DNA [6, 7] and histones [8], as well as regulation of transcription factors [9], RNA interference [10], chromatin remodeling proteins [11], and nuclear architecture [12]. The epigenetic landscape emerging from these alterations enables the storage of more information than is possible using nucleotide sequence alone and represents a powerful force in cellular differentiation and environmental adaptation [13].

In this study, we focus on the recoloring of histones, which plays a central role in controlling gene expression and is strongly related to chromatin folding, as evidenced by the strong correlation with checkerboard patterns in Hi-C contact maps [14]. Thus, the interplay between chromatin organization and spreading of nucleosome modifications (for example methylation of Lysine 9 in Histone 3) is of paramount importance in understanding the dynamical structure and function of chromatin. To explore how spreading is affected by chromatin dynamics, we created a polymer model that employs relative rates of modification and un-modification of nucleosomes. This allowed us to explore the role that chromatin spatial dynamics plays in the formation of modified domains, as well as which regions of chromatin could be associated with heterochromatin.

The intricacies of initiation, spreading, and maintenance of histone modifications currently remain unclear because of the involvement of several factors whose roles have not been quantitatively elucidated. Upon DNA replication, histones are re-distributed in roughly equal proportions to the template and nascent DNA [15]. Subsequently, nascent histones must re-establish the appropriate epigenetic attributes acquired before cell division (‘memory’). Studies on chromatin inactivation have revealed important elements necessary for gene silencing, such as nucleation elements, which are specific DNA segments that bind protein complexes either directly or via RNA interference, and histone-modifying proteins, such as methyltransferases, which covalently modify histone tails [1]. Although molecular identities of these mediators differ across eukaryotic species, certain unifying principles seem to underlie the observed homology between proteins that moderate them in yeasts, humans, mice and flies [16, 17]. Experiments and theoretical studies suggest that a positive-feedback allosteric mechanism is important for the spreading of modifications, whereby the molecular complex binding to the nucleation site has an enhanced propensity to bind to nucleosomes that are already marked by an appropriate enzyme. The bivalent binding is thought to be the basis of the cooperative model of spreading of the modifications [1].

Because of the involvement of several molecular components and inherent stochasticity in the modification process, a variety of mathematical models, which have provided considerable insights into the formation of epigenetic domains and their self-perpetuation, have been proposed [2, 18–27]. Some models may be classified as essentially one-dimensional in which spreading occurs along a lattice representing the chromatin with built-in implicit positive feedback mechanism [22, 23]. Other models consider the possibility of spreading beyond near neighbors of a modified nucleosome, which implicitly accounts for long range contacts [20, 21] along the genomic length. More recently, models that consider the polymeric characteristics of chromatin explicitly [18, 19, 26, 27] or implicitly [21] have been investigated. Here, we introduce a polymer-based model that accounts for the kinetics of modifications, thus coupling chromatin dynamics with stochastic chemical kinetics. The model captures the conformational dynamics of chromatin as well as the biochemical mechanism of spreading, implemented though distinct rules under which the enzyme reactions take place. The aim is to test the extent to which conformational dynamics influence the formation of epigenetic domains, and whether the spreading of epigenetic marks is achieved along the chromatin thread, or is controlled by non-adjacent nucleosomes (see Figure 2). Our model, with a minimal number of parameters, shows that stable epigenetic domains emerge without explicit boundaries when the conformational rearrangement of chromatin dictates the spreading of epigenetic modifications.

**Figure 1:**
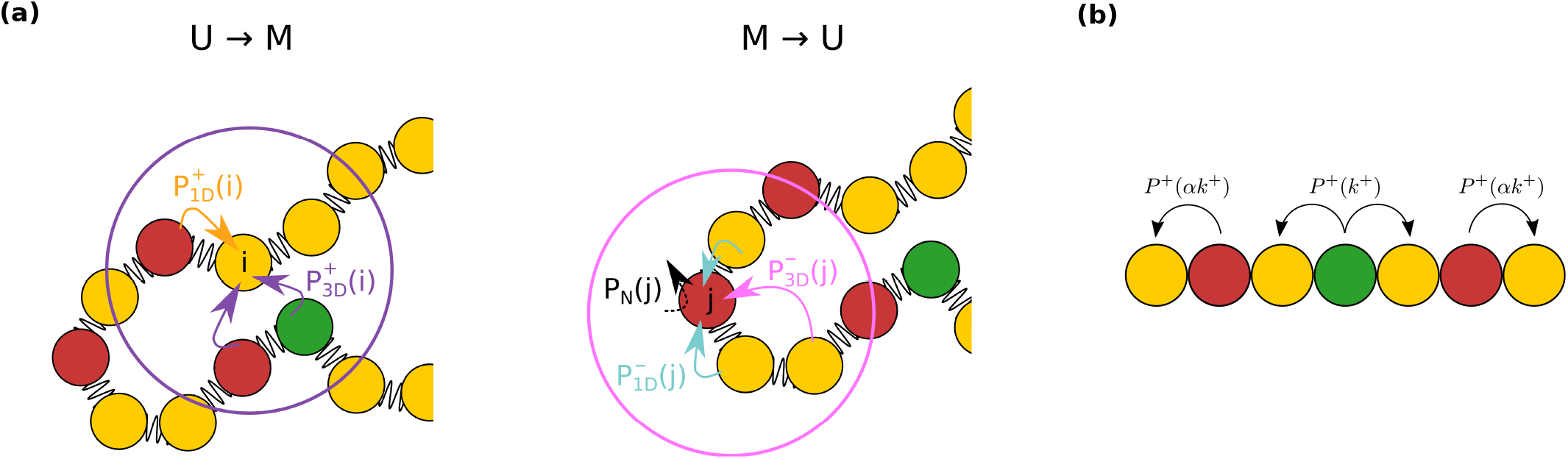
Transitions in the kinetic model. The forward and backward reactions are shown on the left and right panels, respectively. The left panel depicts the enzyme-induced modification, *U* → *M*, at the *i^th^* nucleosome. The right panel sketches the removal of the mark, *M* → *U*, from *j^th^* nucleosome. Modifications to unmarked (U) nucleosomes and removal of marks from the M-type nucleosomes are shown by arrows. Important mediators relevant to the spreading processes are shown in color: the nucleation site (NS) is in green, *U*(*M*) nucleosomes are shown in yellow (red). Some or all elementary transitions are considered simultaneously, representing four separate spreading mechanisms (see Figure 2). The forward reaction occurs if an unmarked nucleosome *i* has a neighbor nucleosome in the modified state with a probability 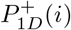 (orange arrow) or with a probability 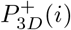 if it is in the spatial vicinity of a nucleation site/modified nucleosome (purple arrow). The backward reaction can always proceed with a probability *P_N_* (*j*), indicated by a black dashed arrow. The *M* → *U* reaction may also depend on the presence of other unmarked nucleosomes. In such cases, neighbor nucleosomes might induce the *M* → *U* transition in nucleosome *j* with a probability 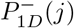 (cyan arrow), while *U* nucleosomes in the spatial vicinity of *j* may induce the reverse reaction with probability, 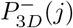 (magenta arrow). **(b)** Asymmetric spreading from the NS (green) and non-NS sites (red). The forward rate for the non-NS sites is scaled by a factor of *α* compared to that for the NS site. Typically, we use *α* ≪ 1 in this study, which means that the ability of spreading of a non-NS site is much smaller than that of the NS site.

**Figure 2:**
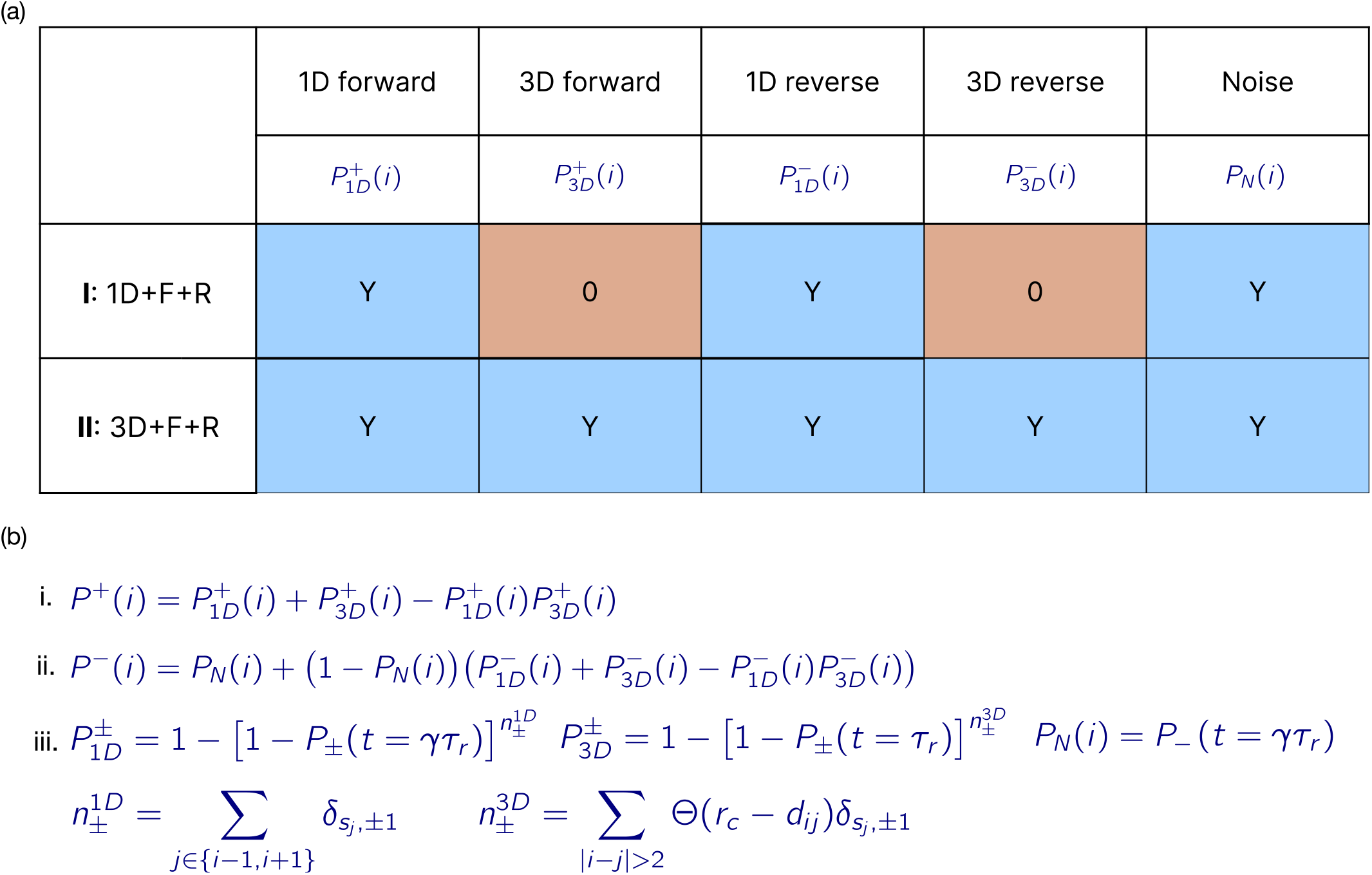
**(a)** Columns represent the elementary reactions and the rows show the schemes implemented in the simulations. The five elementary reactions are: 1D forward, 1D reverse, 3D forward, 3D reverse, and the noise. The noise process, with the associated probability *P_N_*(*i*) where *i* is the nucleosome label, affects only the modified nucleosomes. The two simulation schemes are: **I** (1D+F+R, (1D forward, 1D reverse, and the noise)), and **II** (3D+F+R, (1D forward, 1D reverse, 3D forward, 3D reverse, and the noise)). Y indicates that the probabilities for the five elementary reactions are computed using equation **iii** in **(b)**, and 0 means that the corresponding probability is zero. For each simulation scheme, the cumulative probability for the forward (*P*^+^(*i*)) and reverse (*P*^−^(*i*)) reactions is computed using Equations **(b)i** and **(b)ii**. For scheme **I** (1D+F+R), the probabilities 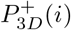 and 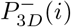 are zero, resulting in 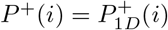 and 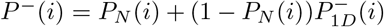, where 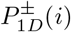 and *P_N_*(*i*) are computed using Equations **(b)iii**. The probability for scheme **II**, which includes all the elementary reactions, is given by *P*^+^(*i*) (Equation **(b)i**) and *P*^−^(*i*) (Equation **(b)ii**). We sometimes report results for special cases of scheme **I** and **II**; scheme **1D** is a special case of Scheme **I** used to assess the spreading due to purely a one dimensional process, whereby scheme **3D** is a special case of scheme **II**, where we don’t include reverse reaction. **(b)** Equations for all the relevant probabilities. (i) and (ii) are the cumulative probability for forward and reverse reaction. (iii) Displays the probabilities for the five elementary reactions. 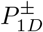 and 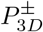 are derived by assuming that each neighbor nucleosome contributes to the forward and reverse transition independently. *t* = *γτ_r_* (*t* = *τ_r_*) is the time at which 1D and noise (3D) spreading is considered. The variables 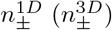 are the number of nearest neighbors (non-neighbor neighbors) whose state is *U* (−1) or M (+1) to *i^th^* nucleosome. Non-neighbor neighbors (*js*) satisfy the criterion |*i* – *j*| ≥ 2 and *r_ij_* < *r_c_* = 1.122*σ*.

We find that the nucleation site acts as a positional signal that facilitates spreading to its surroundings. The bidirectional spreading from the nucleation site was previously explored without considering chromatin conformation explicitly [21] or under conditions in which the formation of an epigenetic domain relies on attractive interactions between similarly-modified loci [19, 27]. As a consequence, partial collapse of chromatin is required for epigenetic spreading. In our model, the cooperative effect of the writing and erasing processes, by which the probability of modification of a locus depends on epigenetic states of the neighbors, provides the sole positive reinforcement for spreading without the need for global conformational change of the chromatin. We show that finite modified domains cannot form without a looping mechanism, which brings nucleosomes that are well-separated along the genome into proximity. Comparison with other mathematical models shows that there are many possible scenarios for modified domain formation, attesting to the complexity of these dynamical processes.

## II. METHODS

### Model

To probe chromatin inactivation, using the modification mechanisms sketched in Figure 1, we used a polymer with *N* = 300 nucleosomes. Each monomer represents 200 base pairs (bps), accounting for one nucleosome and a linker DNA. Hence, the polymer models 60 kb of DNA. The choice of *N* = 300 allows us to efficiently explore the spreading mechanism for a range of parameters. The effects of changing *N* are discussed in the Supplementary Information.

### Energy function

The energy of the chromatin model is a sum of bond stretch (*U_B_*), bond angle (*U_KP_*), and interactions (*U_LJ_*) between the nucleosomes that are separated by at least 3 bonds. A brief description of *U_B_, U_KP_*, and *U_LJ_* follows.

#### Bond stretch and bond angle potentials

The connectivity of the chromatin thread is taken into account using a harmonic potential,

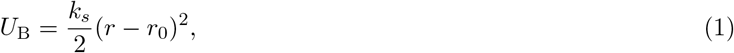

where *r* is the distance between the two consecutive nucleosomes, *r*_0_ is the equilibrium bond length, and *k_s_* is the spring constant. The bond angle is constrained using the Kratky-Porod potential in order to control the stiffness of the chain. We assume that,

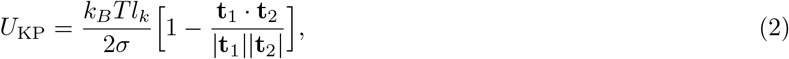

where *k_B_* is the Boltzmann constant, *T* is the temperature, *l_k_*/2 is the intrinsic persistence length (*l_p_*), and *σ* is the effective inter nucleosome distance. The length unit is *σ*. The variables **t_1_** and **t_2_** are bond vectors connecting nucleosomes (*i*, *i* +1) and (*i* + 1, *i* + 2), respectively.

#### Non-bonded potential

We used the Lennard-Jones (LJ) potential,

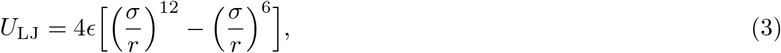

to model interactions between non-bonded loci. In the above equation, *r* is the distance between the nucleosomes, *σ* is roughly the size of the nucleosome, and *∊*, which sets the energy scale, is the strength of the interactions. The LJ interaction is truncated at 3*σ* (*∊* = 0 for *r* > 3*σ*). We assume that the e value is a constant and independent of the modification status of the nucleosomes. Previous studies have assumed that *∊* depends on the epigenetic state [18, 19, 27].

#### Solvent quality

The value of *∊* determines whether the polymer adopts a random coil or is collapsed (see the SI for a detailed discussion). We first simulated chromatin in a good solvent, where both loop formation and epigenetic ergodicity (see the SI) can be readily achieved. In good solvent conditions the *∊* value does not depend on the modification status of the nucleosome. In other words, *∊*_MM_ = *∊*_UU_ = *∊*. To explore other possible scenarios, we also simulated the spreading process, by taking *∊*_MM_ different from *∊*_UU_.

Of particular relevance for epigenetic spreading is the persistence length of *l_p_*, because it affects the kinetics of loop formation, which in turn controls 3D spreading through the looping mechanism. The values of *l_p_* in budding yeast vary considerably [28, 29]. Computational modeling of yeast chromosomes suggest that they can be effectively modeled as flexible polymers [30, 31]. Therefore, we chose *l_p_* = *σ*, thus making our polymer a flexible random coil. We also tested the consequences of varying *l_p_* (see the SI for details).

### Stochastic kinetics for epigenetic modifications

We imagine the *U* state to represent the active chromatin, which transitions to the marked *M* state, inactive chromatin. The modifications are achieved by specific DNA binding enzymes, which we model using suitable first-order rate constants (see below). We assume that changes in *U* at site i are possible only if it is in the vicinity of a nucleation site or another nucleosome is in the *M* state. Since the modification status of the inactivated nucleosomes changes dynamically, the present model may be thought of as a generalization of the Chromatin Copolymer Model (CCM) with static epigenetic attributes introduced previously [32]. Because we consider only two epigenetic states, the nucleosome states can be characterized using Ising spin variables, *s*(*U*) = –1 and *s*(*M*) = 1 [33] except that the spin variables change stochastically with time depending on the instantaneous chromatin conformation. In other words, there could be a cooperative transition in the spin states as chromatin evolves. n the model, positive feedback, mimicking the ‘reader-writer’ model associated with the spreading process, arises because the probability of modifying a nucleosome increases if nucleosomes in the vicinity are already modified. For example, methyltransferases have a reader domain that binds to the previously modified nucleosome, and subsequently write the same modification to another nucleosome [34].

We model the dynamics of enzyme-mediated modifications as a two-state chemical reaction. The nucleosomes undergo a reversible change in their epigenetic states, described by the reaction,

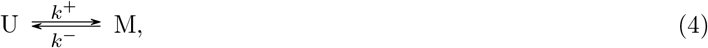

where *k*^+^ and *k*^−^ are forward and backward rates, respectively. The two-state model is a simplification compared to previous studies [18–20, 27], which used three (active, inactive, and unmarked) states. Two-state or effective two-state chemical modifications schemes previously have been used [21, 25].

We calculated the transition probabilities for the 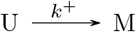 and 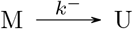 for every nucleosome except for the fixed nucleation site (NS) by solving the rate equations associated with the enzyme-catalyzed reactions. The transition probabilities, *P*^+^(*t*), and *P*^−^(*t*) are evaluated at *τ_r_*, the characteristic relaxation time of the chromatin thread. Using the solutions to Eq 4, we evaluated *P*^+^(*t*) and *P*^−^(*t*), which we write as,

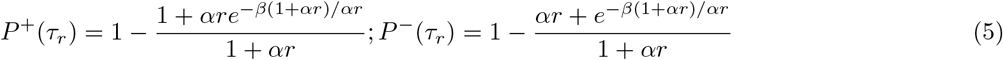

where *r* = *k*^+^/*k*^−^ and *β* = *k*^+^*τ_r_*. We introduce the parameter *α* in Eq 5, which is unity for spreading from the NS (Figure 1). Thus, the probabilities of modification of nucleosomes (Figure 2) depend on *r*, *α*, and the *β* parameter that measures the spreading rate in terms of *τ_r_*. Using this approach, we can relate the probabilities of stochastic events to timescales with physical meaning, as will be detailed below.

### Interpretation of the modification algorithm

The forward reaction is a caricature of chemical modifications of histone 3 (H3) lysine 9 (K9) with methyl groups (me2/3), referred to as H3K9me2/3, a hallmark of inactive chromatin [16]. The modification is catalyzed by Clr4 in *S. pombe* and Suv39h in humans [16, 17]. The reverse reaction takes into account the removal of markers corresponding to histone turnover. In addition, *U* can reverse the state of *M* nucleosomes in their vicinity, thus, introducing the cooperative effect in the reverse reaction. The latter process may mimic the activity of enzymes that remove histone modifications (for example, Epe1 in *S.pombe* [35]).

At each nucleosome *i*, we envision five transitions (Figure 1) that could lead to modifications based on the kinetics in Eq 5. These transitions are attempted at every time step of the evolution of the chromatin. They are (shown schematically in Figure 1 and their equations are shown in Figure 2):

1. 1D: *U* → *M* transition may occur only if at least one nucleosome at *i* ± 1 is in the *M* state.
2. 3D: *U* → *M* transition may occur only if there is a minimum of one nucleosome, *j*, in state *M* that satisfies the criteria, |*j* – *i*| ≥ 2 and the distance *r_ij_* to *i^th^* nucleosome is less than the threshold, *r_c_*.
3. Noise: *M* → *U* transition could occur at any nucleosome independent of the identity of all others.
4. 1D: *M* → *U* transition may occur only if least one nucleosome at *i* ± 1 is in the *U* state.
5. 3D: *M* → *U* transition may occur only if there is a minimum of one nucleosome, *j*, satisfying |*j* – *i*| ≥ 2 and *r_ij_* < *r_c_*.

Because we are mostly interested in epigenetic spreading, we do not consider the spontaneous stochastic *U* → *M* transition (the analog of the third transition listed above).

Nucleosomes in the *M* state could probabilistically spread the mark, which may be thought of as caricatures of the biological mechanisms (reader-writer or writing only given in step 3 in the scheme given above). The complicated processes involved in enzyme-induced marking or unmarking are subsumed in single rate constants and the probabilities of the modifications. We introduce a parameter *α* that differentiates between the spreading rates from the NS (*αk*^+^ where *α* is unity), and nucleosomes that are modified through the kinetic scheme listed above (*αk*^+^ with *α* less than unity, as illustrated schematically in Figure 2). The *M* → *U* reaction, which erases or removes the mark, occurs at a rate *k*^−^ (steps (4) and (5) described above). The value of *k*^−^ is set to be a fraction of the forward rate, *k*^+^ > *k*^−^. Modification in step 3, at the rate *k*^−^, is associated with enzyme turnover, occurring stochastically, and is sometimes referred to as noise. The noise term is always present and does not depend on the states of the neighbors.

At time step, *t* = 0, all the nucleosomes are unmarked, except for the nucleation site (shown in green in Figure 1), chosen arbitrarily to be in the middle of the polymer, although we investigated the effects of changing its location (discussed in the SI). Spreading starts at the NS, which enables bidirectional spreading along the nucleosomes, emanating outward from its location. A spreading trajectory is generated by stochastic changes in the nucleosome states, as outlined above in the five steps. The probabilities associated with such changes are intimately coupled to the chromatin dynamics, as we explain below.

### Epigenetic clock

The time that determines 3D spreading is associated with the lifetime of contact between two nucleosomes *i* and *j* with |*i*–*j*| ≥ 2. There is a spectrum of lifetimes associated with loop formation times depending on |*i* – *j* |. Our goal is to elucidate how the chromatin dynamics are coupled to the enzyme-catalyzed reactions that modify the *U* state or erases the mark in the *M* state. Therefore, to set the overall time scale (time unit of the ‘epigenetic clock’), we chose the polymer relaxation time, *τ_r_*, calculated from the time-dependent decay of the structure factor *F*(*q, t*) (see the SI), evaluated at the wave vector *q* = 2*π/r_c_*. We show in Figure S5 that *τ_r_* is a reasonable surrogate for the mean contact lifetimes during which modifications could occur by the looping mechanism. The dependence of the spreading on *l_p_* and chain length *N* are described in the SI. Note that in contrast to previous polymer models, there is an explicit connection between chromatin relaxation time (*τ_r_*) and the dynamics of spreading. We set *τ_r_* (≈ 240s) as the unit of time, and all other times are measured relative to *τ_r_*.

### Fast and slow spreading

We consider the limiting case when epigenetic spreading occurs rapidly compared to *τ_r_*. In the fast spreading (FS) limit, *k*^+^*τ_r_* = *β* ≫ 1, which implies that modification occurs on time scales that are much shorter than *τ_r_*, the chromatin relaxation time. In this limit, spreading occurs predominantly by linear or 1D mechanism with the looping mechanism playing a less significant role. To achieve this limit in the simulations, we introduce a parameter *γ* (Figure 2) only in the 1D dominated FS to ensure that the 3D contribution to epigenetic spreading is minimized. The precise value of *γ* is irrelevant as long as *P*^+^(*t* = *γτ_r_*) is large. In the FS limit, this is achieved by choosing *βγ* > *C*(≈ 3 ~ 4).

The limit of slow-spreading (SS) is achieved by choosing *k*^+^*τ_r_* = *β* ≪ 1. In the SS limit, chromatin undergoes multiple cycles of relaxation, which implies that nucleosomes separated by large genomic distance a have substantial probability of being in proximity for spreading to occur by the looping mechanism. With our choice of *β* = 0.01, we find numerically that spreading by the 1D mechanism is not as important. This implies that in the SS limit the value of *γ* has no bearing on the results of the simulations. Thus, by changing *β* one can examine a range of possibilities for epigenetic spreading.

### Feedback in the model

Our model explicitly accounts for feedback, since the probability of modification of a specific nucleosome is enhanced if there is a neighboring nucleosome in proximity that is in the *M* state. Consider the expression for 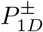 in Figure 2. The allowed values of 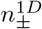 are 0, 1, and 2. As 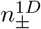 changes from 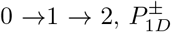 increases. In other words, modification probability increases if there is a neighbor in the *M* state. Similarly, as 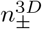 increases the value of 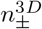 increases as does the probability of modification (see step (iii) in Figure 2). Thus, feedback is automatically incorporated by choosing the modification probabilities in a natural way.

### Transition probabilities

We implement the coupling between chromatin evolution and stochastic modification of the state of the nucleosomes using the scheme in Figure 2. This requires calculation of the probabilities of modifications, 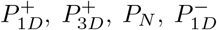, and 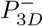. The expressions for these are given in Eq. (iii) in Figure 2. The cross-terms in Eq. (iii) ensure that only one nucleosome *i* is subject to modification in the forward transition or the reverse transition at each step. Scheme **II** mimics the important biological processes that modify the *U* state by 1D and 3D looping mechanisms.

The dynamics of spreading depends on *r* = *k*^+^/*k*^−^, *β*, and *α*. In principle, *r* is measurable because *k*^+^ and *k*^−^ are enzyme catalyzed reactions that methylate or acetylate specific reactions in histones. Changes in the parameter *β* which measures the spreading rate relative to the chromatin relaxation rate, allows us to probe the limit ranging from FS to SS.

In contrast to all previous computational models, the *α* parameter plays a special role in our simulations. In particular, *α*, which is one for the nucleation site (NS) (shown in green in the Figure 1), ensures that spreading from NS is more probable regardless of whether the mechanism is purely 1D or is driven by 3D looping. By having the NS being permanently modified, we show that stable finite domains form without having to introduce any boundary element.

### Implementation

We start from an equilibrated conformation of the chromatin polymer with all the nucleosomes in the *U* state at *t* = 0. The simulations were also repeated with all nucleosomes in the *M* states. The steady-state behavior is independent of the initial conditions, which is assured because the chromatin is epigenetically ergodic (see SI for details). In addition, the simulations were performed for times that exceed *τ_r_* by several orders of magnitude, which implies that the sampling of the conformations is more than adequate.

The conformation of the polymer is evolved by integrating the Langevin equations by choosing a suitable time step 6 (described in Section **1** in the SI). At each time step, marking or unmarking of all the nucleosomes is attempted with probabilities given in Figure 2. We implemented the dynamics, described mathematically in Figure 2, using an in-house computational code. A graphical representation of the modification algorithm is provided in Figure S1 in the SI. A pictorial representation of the simulated trajectories for fast and slow-spreading limits are shown in Figure 3, and in supporting movies M1 and M2.

**Figure 3:**
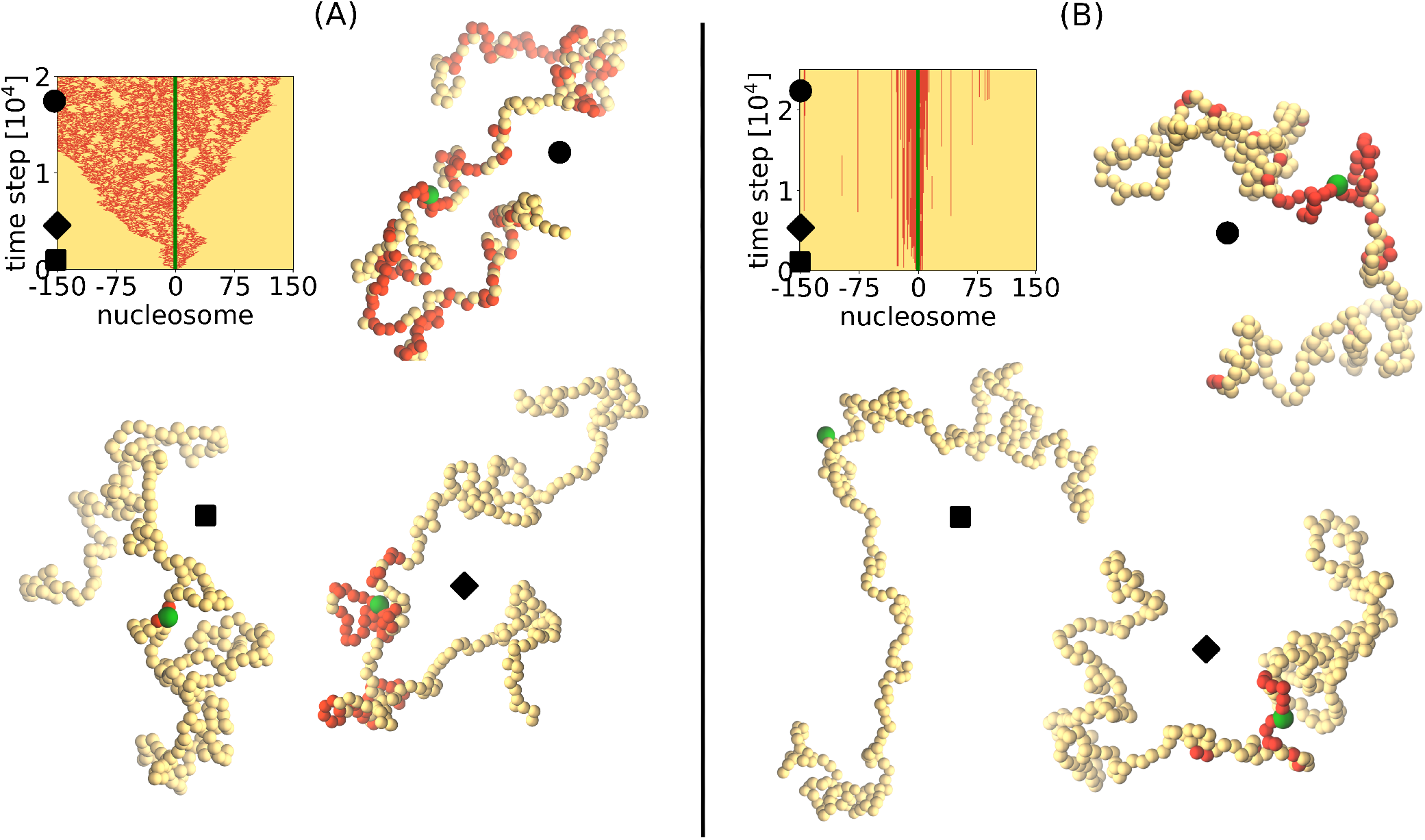
Visualization of single trajectories in the fast (A) and slow (B) spreading regimes. In the former case, the spreading occurs fast and independently of chromatin configuration, while in the latter case, the pattern of spreading may be determined by the dynamic rearrangement of the chromatin, as explained in the main text. The kymographs show the epigenetic identity of each nucleosome (marked/M/red, unmarked/U/yellow, nucleation site/green) for a single trajectory, in both (A) and (B). Three distinct simulation time steps are chosen, marked with a symbol on the kymograph, and we extract polymer conformations with epigenetic identities of the nucleosomes in order to graphically represent them. For clarity, the size of the nucleation site is enhanced. Accompanying movies M1 and M2 are available online.

### Global epigenetic state

We use simple measures to characterize the global epigenetic state of chromatin as well as the modification state of the individual nucleosomes for both the FS and SS scenarios. By examining the dynamics in detail we can quantitatively assess the contributions from 1D and 3D spreading separately for the two extreme caess. We determine the epigenetic state of chromatin using,

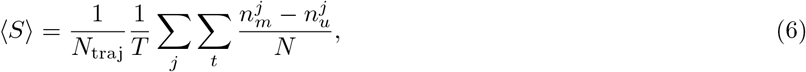

where 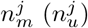 is the number of modified (unmodified) nucleosomes at time *t* in the *j^th^* trajectory, and *N* is the total number of nucleosomes. The quantity 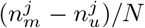 is averaged over time *T* and *N*_traj_. Unless otherwise stated, all the relevant quantities are calculated using *N*_traj_ = 10, where each trajectory starts with a different initial chromatin conformation.

### Average epigenetic state of the nucleosomes

We also determined the average, 〈*s_i_*〉, of each nucleosome, which is calculated using,

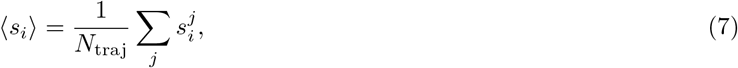

where 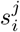 is the epigenetic state of locus *i* in trajectory *j*.

## III. RESULTS

### Fast spreading results in an abrupt switch in the epigenetic state

The interplay between the three parameters, *k*^+^, *k*^−^, and *α*, determines the global epigenetic state. Let us first consider the FS case, where the spreading rate (*k*^+^) is much faster than the polymer relaxation rate 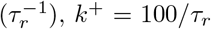. The *k*^−^ value is a fraction of the forward rate, which means the reversal of *M* occurs at a slower rate compared to modification of the *U* state. We follow the modification status of all the nucleosomes over the course of the simulation time in order to assess if the chromatin is predominantly in the global active (unmarked with 〈*S*〉 ≈ −1) or inactive (marked with 〈*S*〉 ≈ 1) state.

Plots of the global epigenetic state and the nucleosome-dependent state in Figure 4 allow us to draw a few pertinent conclusions. (i) Modifications are highly improbable if *k*^+^/*k*^−^ is less than a certain value (≈ 10) regardless of the value of *α*. The chromatin remains in the global *U* state with 〈*S*〉 ≈ −1, regardless of whether the modifications occurs by Scheme **I** or Scheme **II** (see the left panels in Figure 2). In general, *αk*^+^ has to exceed *k*^−^ for spreading to occur. At higher values of *k*^+^/*k*^−^ there is a global *U* → *M* transition as *α* is increased. (ii) The transition from a predominantly active to predominantly inactive state occurs over a narrow range of *k*^+^/*k*^−^, and *α*. For instance, for both mechanisms, **I** and **II**, the switch from 〈*S*〉 < 0 to 〈*S*〉 > 0 occurs over a narrow range of *α*, as is evident from the left panels in Figure 2. The critical point in the switch from active to inactive state is determined by *k*^+^/*k*^−^ (Figure 2 and Figure S8). The transition hinges on the competition between the forward (favors spreading) and the reverse reaction (inhibits spreading) distally from the NS, and is controlled by *αk*^+^ ≈ *k*^−^. Under this condition, the modification results in the concurrent formation of similar-sized patches containing *U* and *M* nucleosomes, resulting in 〈*S*〉 ≈ 0. This is most evident in spreading by mechanism **II** (lower left panel in Figure 2), showing a switch from active state (yellow color, 〈*S*〉 < 0) to the inactive state (red color, 〈*S*〉 > 0) through the mixed state (white squares, 〈*S*〉 ≈ 0). The delicate state of the domain, depending on a narrow *α*(*k*^+^/*k*^−^) range produces an epigenetic switch, reminiscent of gene control expression in cells, whereby cellular fate depends on the relative concentrations of the epigenetic regulators. Shifting the balance in one direction or the other could result in dramatically different outcomes in terms of gene expression patterns [36]. (iii) If *αk*^+^/*k*^−^ < 1 then the reverse reaction occurs predominantly by random histone turnover (‘noise’ in the list of transitions) that does not depend on the epigenetic identity of neighbors. On the other hand, if *αk*^+^/*k*^−^ > 1 the reverse reaction is facilitated (Figure 2 and Figure S8) due to positive feedback, which pushes the domain state towards the global *U* state. To mitigate this effect, stronger enhancement of the forward reaction is needed to achieve silencing levels comparable to the reverse noise dominated mechanism. (iv) The middle panels in Figure 2 show ensemble-averaged (Eq. 7) spin states of each nucleosome as a function of time. First, spreading occurs bi-directionally along the genomic length from the NS. Second, a comparison of the top and bottom panels shows the negligible difference between the **I** and **II** mechanisms. This is reasonable because before the polymer can relax, the modification would have already occurred with high probability implying that chromatin dynamics do not play a significant role. (v) In the fast-spreading limit, domains form without limit, as shown in the right panels of Figure 4. The fraction of modification in time per nucleosome, given by Eq. 6, outlines the temporal stability of the domain without bound. The lower *f_im_* values at chain ends are finite-size effects (see SI and Figure S13).

**Figure 4:**
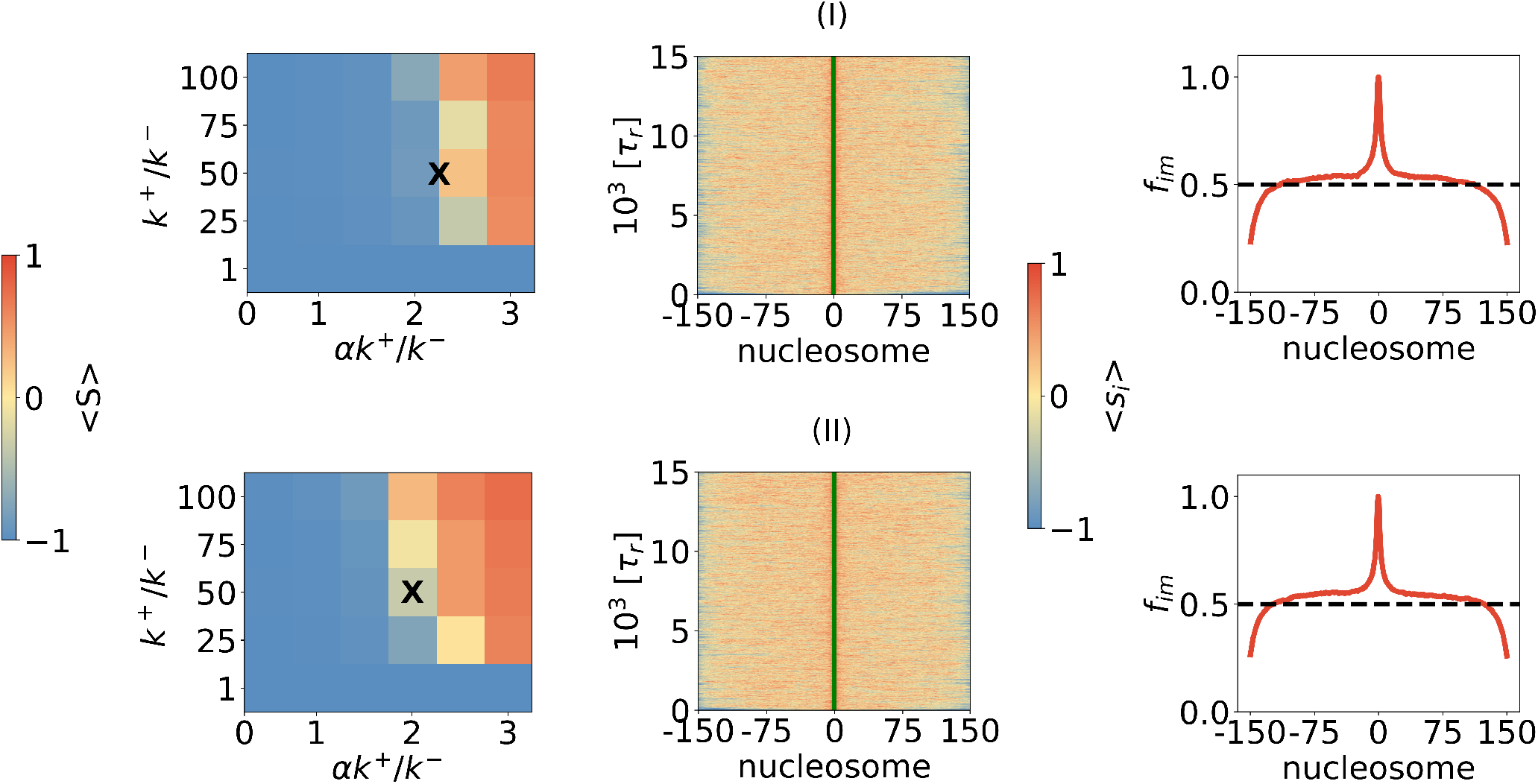
Epigenetic switching depends on modulation of the forward vs back rate distally from the nucleation site. Left panels: Overall epigenetic state, characterized by 〈*S*〉 (Eq. 6). Middle panels: Average epigenetic state of each nucleosome 〈*s_i_*〉 (Eq. 7) with *α* = 0.0446 for Scheme **I** and *α* = 0:04 for Scheme II, marked by X in left panels. Right panels: Fraction of time each locus spends in the modified state (Eq 6 in SI). The parameters are chosen such that the global average 〈*S*〉 ≈ 0. In all the panels, *k*^+^ = 100/*τ_r_* the total length of the simulations is 15; 000*τ_r_*, and *k*^−^ = *k*^+^=50. Vertical green line is the position of the nucleation site.

### Topology-driven domain formation

We next explored the slow-spreading case, mimicked by choosing *k*^+^ = 0.01/*τ_r_*. In this limit, the chromatin polymer could form the allowed contacts through the looping process multiple times on the time scale ≈ 1/*k*^+^, which would allow for 3D as well as 1D spreading. Consequently, the inactivation profile would be determined by the contact probability between the nucleosomes separated by | *i* – *j*| ≥ 2 (Figure S5).

These expectations are tested by computing the epigenetic states of the nucleosomes as well as the associated inactivation profiles. We fixed *k*^+^ and varied *k*^−^ to improve the odds of spreading, deciding finally on the choice *k*^−^ = *k*^+^/10, 000. For the chosen parameters, domain formation by the linear mechanism does not occur (Figure 5). This is because the probability of linear spreading to neighboring nucleosomes, even away from the NS, is extremely low (≈ 6 o 10^−5^) at each time step. Thus, we surmise that spreading must occur exclusively through the formation of 3D contacts. This is borne out in the bottom panels in Figure 5, which show clearly that the 3D spreading mechanisms result in the formation of stable domains around the NS. It is worth noting that stable domain formation (≈ 60 nucleosomes centered around the NS) does not require collapse [18] or partial collapse [19] of the chromatin polymer.

**Figure 5:**
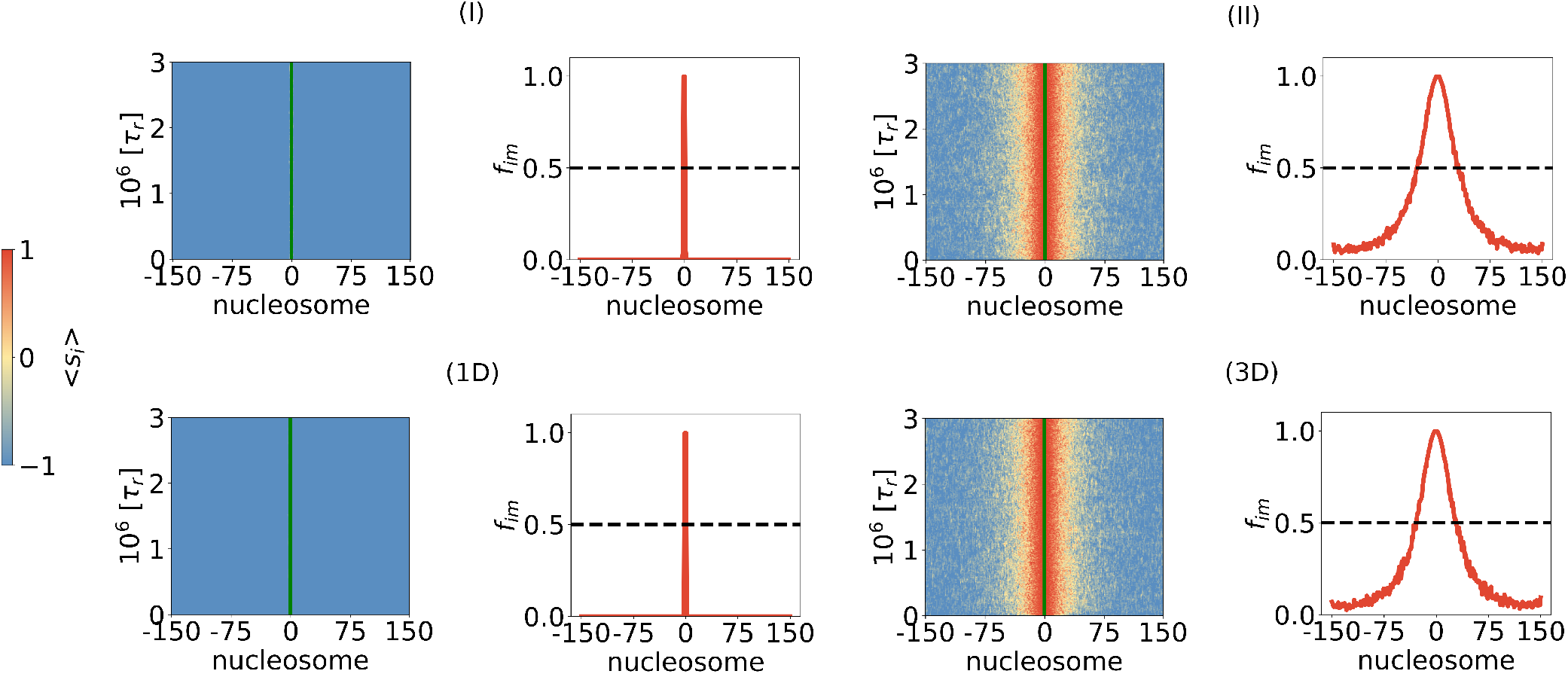
Formation of epigenetic domain determined by slow spreading kinetics with *k*^+^ = 0.01/*τ_r_*. Left panels show average (over an ensemble of trajectories) states of nucleosomes as a function of time. The panels on the right show mean values (Eq.6) of the spin states of the nucleosomes. These are the profiles that show the extent of inactive domain formation. The value of *k*^+^/*k*^−^ = 10,000 in all simulations. The label ID is a mechanism obtained as the limit of **I** in which the modification probability is calculated using *P*^+^(*i*) + *P*^−^(*i*) with 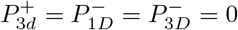. Similarly, 3D represents a process derived from **II** with 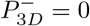. The value of *α*(1*D*) = 6.3·10^−5^, *α*(**I**) = 2.23 · 10^−4^, *α*(3*D*) = 4.5·10^−5^, *α*(**II**) = 2·10^−4^.

The spreading profiles in the bottom right panels of Figure 5 show that the peak in the *f_im_* profile is localized around the NS. The boundary (or the interface) between the active and inactive domains is relatively soft [37], indicated by a continuous decrease of *f_im_*, rather than by a step-like drop to 0, which would occur if the boundary were sharp. Thus, the percentage of inactive loci in the interface region between the active and inactive nucleosomes could indicate whether the boundary is efficient in preventing modified nucleosomes from spreading distally of its position.

An important finding in our work, with potential biological importance, is that even without an explicit boundary element, the epigenetic domain size is finite, and is localized around the NS. The distribution of modifications around the NS is governed by the contact probability of the NS with surrounding residues, which depends on the chromatin conformational dynamics, and less so on the reverse reaction. This is evident by our finding that the inactivation profiles for **II** and (3D) scheme, which disallows the reverse reaction, are similar (bottom panels of Figure 5). Furthermore, the shape of the epigenetic domains shown in the bottom panels of Figure 5 and the boundary formation is due to the asymmetry in the spreading rates from the NS (*k*^+^) and the modified nucleosomes (*αk*^+^ with *α* less than unity). This asymmetry is required for the formation of discrete domains without explicit boundary elements to halt the spreading process, and points to the importance of the NS that we explore further (see below).

### Inactive domains cannot form without the NS

The results in Figure 5 show that the presence of the NS results in finite-sized inactive domain formation. We also find that *f_im_*, even in the fast-spreading case 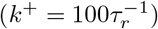, is far less than 0.5 in the absence of the NS (Figure S15). The kymographs for all four (1D, **I**, 3D, and **II**) mechanisms, equivalent to the ones in Figures 5 and S7, show that the nucleosomes are predominantly in the unmarked state. Another striking feature that characterizes the no-NS situation is the increased homogeneity in the distribution of the marked states along the chromatin. This is vividly shown in the right panels in Figure S15, showing flat inactivation profiles with *f_im_* < 0.5. The reverse reaction (*M* → *U*) shifts the inactivation percentages for the **II** mechanism well below those of others (see the bottom right panel in Figure S15). Taken together, these results establish that no NS implies that a finite-sized inactive domain with high probability cannot exist.

### DNA replication and the NS

Next, we address the role of the NS in a pre-formed epigenetic domain and the impact of DNA replication. In our model, the NS is the only element whose epigenetic identity is unchanged, and functions as a reservoir for modifications. We first performed simulations with *α*-values that result in a halfsilenced domain, 〈*S*〉 ≈ 0. At *t* = 15, 060*τ_r_* (*t* = 301, 205*τ_r_* for slow-spreading) we removed the NS by changing it to a nucleosome in the *M* state. With this alteration, the identity of the *M* state could change stochastically as the chromatin evolves, unlike the NS. At *t* = 30,120*τ_r_* (*t* = 602, 410*τ_r_* for slow-spreading), we mimic DNA replication by randomly assigning the active state to half of the nucleosomes. Subsequently, DNA replication is repeated every 150.6*τ_r_*. Although the replication time is arbitrary, it turns out that it corresponds to ≈ 10 hours because *τ_r_* ≈240s (see the Discussion section). Note that the end-to-end distance of the polymer, which is the lowest degree of freedom of the chromatin, relaxes at about 602.4*τ_r_*, indicating that the DNA replication time (150.6*τ_r_*) is shorter than the polymer relaxation time. By following the epigenetic identities of all the nucleosomes, we calculate the inactivation profile of the chromatin from the portion of the trajectory after removing the NS. We define epigenetic domain to exist in the region above the level *f_im_* > 0.5.

Figure 6 shows the results for scheme **II** for fast (Figure 6A) and slow (Figure 6B) spreading, and the scheme 3D for fast (Figure 6C) and slow (Figure 6D) spreading. On the left panels of all four subfigures in Figure 6, the first part of the trajectory where the NS is present, is identical to the kymographs in the right panels of Figures 2 and S7. Upon removal of the NS, for Scheme **II** with fast-spreading, the domain is maintained after the nucleation site is deleted (red curve in the Figure 6A). However, the level of *f_im_* falls below 0.5 when DNA replication starts, suggesting that such a domain is destabilized with DNA replication acting as a perturbation. Notwithstanding, if the backward rate is enforced to be zero 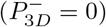, the domain is maintained during DNA replication (Figure 6C). This can be expected as such system, lacking positive-feedback *M* → *U* transitions, is naturally more resistant to maintaining a marked *M* state. In the limit of slow-spreading, shown in subfigures B and D, the finite-sized domain cannot be maintained once the nucleation site is deleted (Figure 6B,D). Stable, bounded domains that are maintained during DNA replication would only occur if the looping mechanism for modification is allowed and the NS does not change. Thus, the results in Figures 6A and 6B show that the NS is required for domain maintenance if DNA replication is permitted. Experimental evidence on *S. pombe* [38] does suggest that some nucleation events needed for heterochromatin formation are triggered at each cell cycle, indicating that maintenance requires the nucleation event.

**Figure 6:**
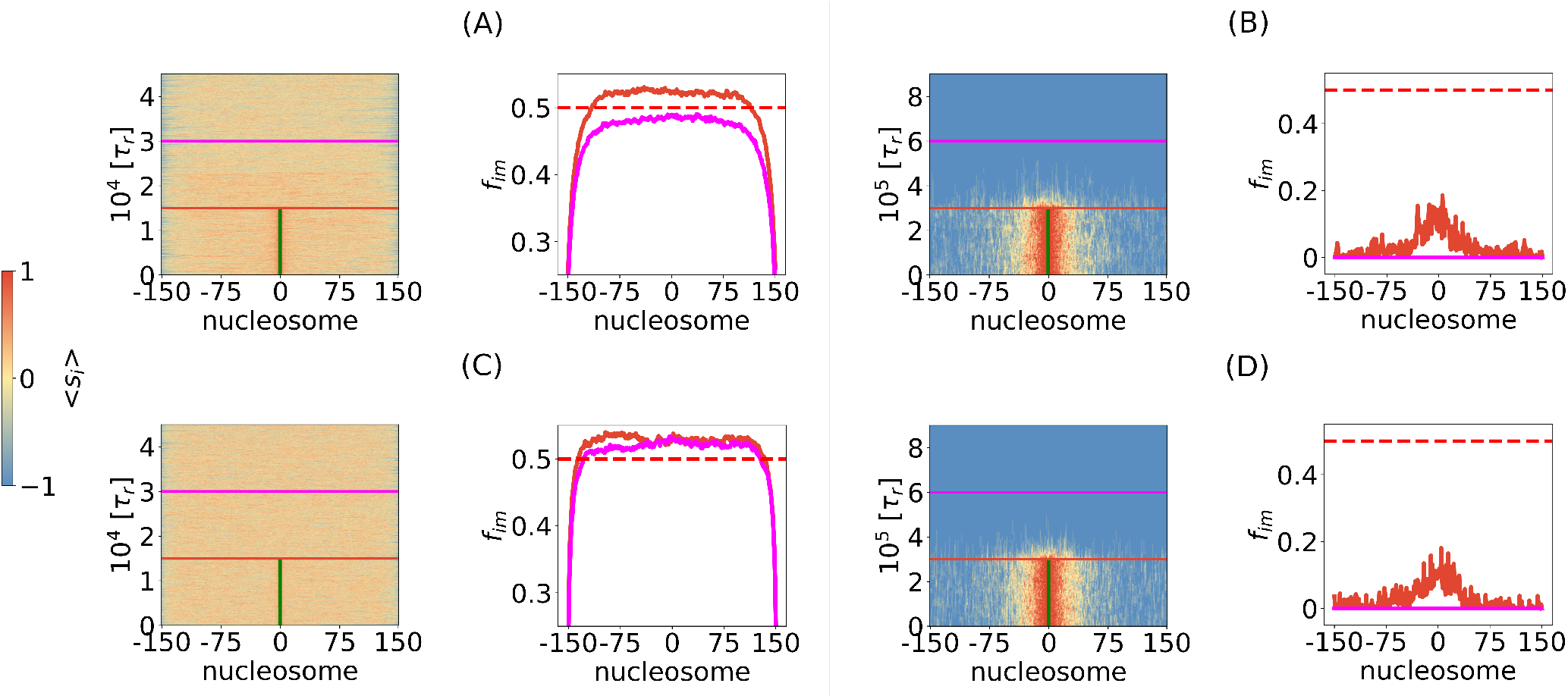
Role of nucleation in establishing and maintenance of epigenetic spreading. (A) and (B) correspond to Scheme **II** in the fast (A) and slow (B) spreading regime. (C) and (D) correspond to a special case of Scheme **II**, denoted as 3D (explained in captions of Figure 2 and 5), also presented in the fast (C) and slow (D) spreading regime. In each of the four subfigureβ, the left panel is the state of each nucleosome in a trajectory, 〈*s_i_*〉 (Eq 7). The trajectory is divided into three parts with distinct simulation conditions, whose transition is marked by two horizontal lines. Initially, the simulation starts with the NS present. Subsequently, the NS is removed (marked by red horizontal line). After removing the NS, a new steady state is achieved. Thereafter, we introduoe DNA replication (marked by magenta hcrizontal line) by stochastically replacing approximately 50% of *M* marks with *U* marks, mimicking equal redistribution of parent histonea to both strands upon replication. Replications are repeated every 150.6*τ_r_*. Right panels in all subfigures indicate the potential presence of epigenetic domains, observed if the fraction of trajectory in marked state for each locus (Eq 6 in SI) is > 0.5. Horizontal dotted represents the 50% mark. For all fast spreading simulations, *k*^+^/*k*^−^ = 50, *k*^+^ = 100/*τ_r_*, while for slow spreading simulations *k*^+^/*k*^−^ = 10,000 and *k*^+^ = 0.01/*τ_r_*.

### Finite domains in an initially condensed or partially condensed chromatin

The result presented so far were obtained by setting *∊*_MM_ = *∊*_UU_ = 0.1*k*_B_*T*. This choice places the chromatin thread in a good solvent, which could mimic the behavior in fission yeast [39]. However, it is believed that in eukaryotes heterochromatin is denser than euchromatin. In the context of our model, it would imply that the effective interaction *∊*_MM_ > *∊*_UU_. To account for the higher chromatin density, we set the interaction strength *∊*_MM_ = 3*∊*_UU_, which implies that due to dynamical changes in the epigenetic landscape, the effective interactions between nucleosomes are modified as in the previous polymer-based studies [18, 19, 40]. We set *∊*_UM_ to the geometric mean, 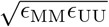. Since linear spreading does not depend on solvent quality, we only explored Scheme **II**, where looping determines the outcome of the spreading process. We simulated the SS case (*k*^+^*τ_r_* = *β* = 0.01).

To fully explore the range of solvent conditions, we considered two cases. In both, we chose *∊*_MM_ = 3*∊*_UU_. Let us first consider *∊*_UU_ = 0.1*k*_B_*T* and *∊*_MM_ = 0.3*k*_B_*T*. For a homopolymer, *∊*_MM_ = 0.3*k*_B_*T* is close to but slightly below the Θ point (see Fig. S3 in the SI) whereas for *∊*_UU_ = 0.1*k*_B_*T* chromatin behaves as disordered random coil. With this particular choice, we find (upper panels in Figure 7) that the results are qualitatively similar to the case with *∊*_MM_ = *∊*_UU_ = 0.1*k*_B_*T* (Figure 5). In particular, the domain size is bounded, as discovered in experiments on mouse stem cells and fibroblasts [37]. This is outlined both by the pattern formed around the nucleation site on the kymograph (upper left panel), as well as in the activity profile of the domain (upper right panel). Since the epigenetic spreading in this regime is mostly due to contacts between the NS and spatially nearby nucleosomes, spreading is localized.

**Figure 7:**
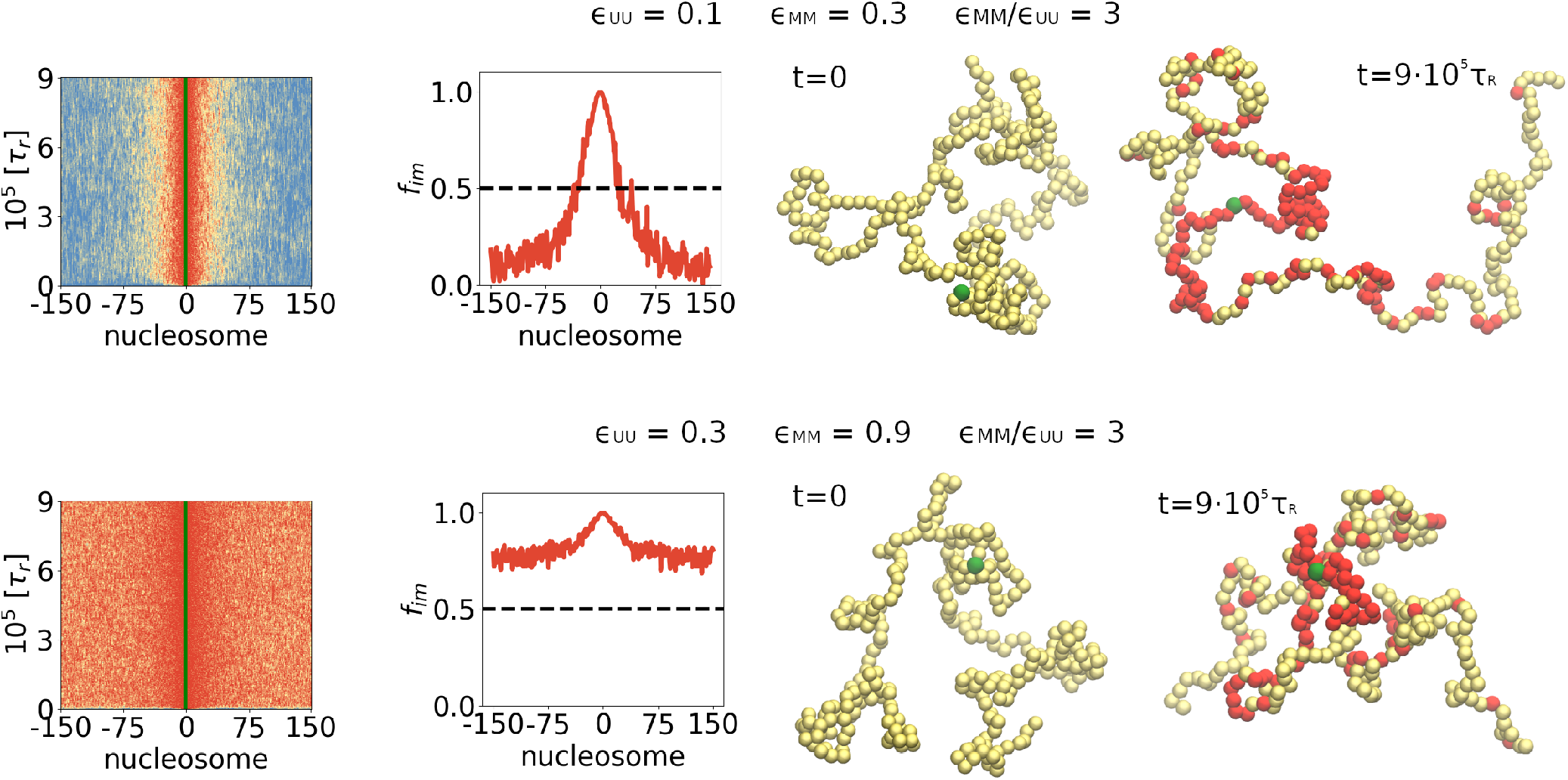
Effect of solvent quality on the spreading of epigenetic modifications using Scheme **II**, in which looping affects the spreading process. The(*∊*_UU_) and *∊*_MM_ values are given on top of each panel. For all simulations, *k*^+^/*k*^−^ = 10,000 and *k^+^* = 0.01/*τ_r_*. The initial and the final conformations of the chromatin are displayed on the right.

The *∊*_MM_ = 3e*∊*_UU_ with *∊*_UU_ = 0.1*k*_B_*T* corresponds to a mixed case in which a fraction of modified nucleosomes experience moderately poor solvent conditions whereas the *U* nucleosomes are in good solvent (see Fig.S3 in the SI). We then investigated the case when the modified nucleosomes are in bad solvent conditions and unmodified nucleosomes are close to but slightly below the Θ condition by choosing *∊*_MM_ = 3*∊*_UU_ = 0.9*k*_B_*T* while keeping the ratio *∊*_MM_/*∊*_UU_ = 3 as before. The results in the lower panels in Figure 7 are dramatically different from those in the upper panels. Spreading of the marks is centered around the NS as before. However, modification occurs without bound with the probability of the *i^th^* nucleosome to be in the M state is ≈ 0.7 as the genomic length increases or decreases from the NS. In contrast to the results in Figure 4, which shows that probabilities of being in the M or U state away from the NS are ≈ 0.5 (except for the ends) the results in the lower panels demonstrate a substantially higher probability of being in the M state. Taken together the results in Figure 7 suggest that *∊*_MM_/*∊*_UU_ is not as important as the solvent quality for predicting the outcome of the spreading dynamics. Although *∊*_MM_/*∊*_UU_ = 3 in Figure 7, the initial conformation of the chromatin is a disordered coil in the upper panel whereas in the lower panels it is more compact. The final epigenetic state of chromatin can be anticipated by the solvent quality of that the chromatin is in at the initial state, especially in the SS limit, where spreading occurs predominantly by the 3D mechanism.

## IV. DISCUSSION

We developed a minimal polymer model to explore different scenarios for epigenetic domain formation with a focus on the coupling between chromatin dynamics and stochastic switching in individual nucleosome states. We find that the interplay of the structural relaxation rate and the modification rates determine the efficacy of the spreading process. However, by setting our work in the context of the modeling literature on epigenetic spreading, we conclude that multiple scenarios for heterochromatin spreading are possible because the domain formation is an interplay of several time, length, and energetic scales. A similar picture emerges from experimental studies, encompassing different organisms and epigenetic modifications.

The main findings in our work are:

- In the limit of fast-spreading, corresponding to the *U* → *M* modification rate much greater than the relaxation rate of the polymer, the spreading occurs predominantly linearly. If the ratio between forward and reverse rate *k*^+^ /*k*^−^ is sufficiently large, unbounded heterochromatin domain forms rapidly (time scales on the order of ≈ 100*τ_r_* as shown in Figure S17 in the SI). In the slow-spreading limit, with the *U* → *M* modification rate that is much less than the relaxation rate of the polymer, finite domains are established on the time scales of ≈ 10, 000*τ_r_*. The domain size is ~ 60 nucleosomes or roughly 12kbs. The width of the interface between the active and inactive domains is soft involving a small number of nucleosomes (Figure 5).
- The presence of the nucleation site is essential for the formation of the modified domains in both the fast and slow-spreading scenarios. In the fast-spreading limit, if a domain has been established and then NS is removed, the domain remains stable. In slow-spreading, this is not the case, an already established domain cannot be maintained if NS is removed. It should be noted that there is evidence for the NS in experiments [37]. Somewhat surprisingly, DNA replication preserves the domains, which might have implications for heritability.
- An interesting finding is that, in the limit of slow-spreading, finite modified domains without boundary elements that are known to stop the spreading process. Finite domains form by a mechanism **II** that predominantly involves 3D chromatin looping. Finite domains form by mechanism II predominantly involving 3D chromatin looping.
- A major determinant of spreading is the solvent quality of the initial (fully unmarked) epigenetic state, rather than the relative strength of the *U* – *U* and *M* – *M* interactions. If the initial epigenetic state of the chromatin is a coil, contacts rarely form, resulting in localized spreading around the nucleation site, which leads to a discrete bounded domain (upper panel in Fig. 7). If initially chromatin is in just below the Θ-point we find that stable finite domains form (upper panel in Fig. 7) even if the interactions between the modified nucleosomes are attractive. On the other hand, if the initial chromatin state is such that it is partially condensed, we find that epigenetic domains form without bound (lower panel in Fig. 7). Because propagation of heterochromatin without bound is biologically untenable, our results suggest that either there ought to be multiple boundary elements that stop the spreading of epigenetic marks or the environmental conditions for the unmodified nucleosomes should poise them close to the Θ conditions.

### Comparisons to previous studies

Several models, which are based on stochastic kinetics to mimic the process of spreading, have been proposed. These studies are based on models in which spreading occurs linearly [25] or through a combination of linear mechanism and implicit long-range effects [20–22]. These studies have insights into epigenetic spreading and inheritability but did not explicitly consider the polymeric nature of chromatin. Our approach, accounting explicitly for the polymer features of chromatin together with stochastic changes in the epigenetic states of the nucleus, is closest to three previous insightful studies [18, 19, 40] but differs in important ways.

(i) In the previous studies [18, 19] the interaction between the nucleosomes changes dynamically as the chromatin evolves. The variations in the strength of the epigenetically modified interactions between nucleosomes of the same state are dynamically altered (Brownian [18] or Monte Carlo dynamics [19]). Thus, the coexistence between silenced and active states is a consequence of the emergent asymmetry in the interaction strengths between nucleosomes of like and unlike epigenetic identity [18, 19]. In contrast, in our model even with *∊*_MM_ = *∊*_UU_, (with the same interaction between the marked and unmarked nucleosomes) finite domains form. The reason is that domain formation and switches from modified to unmodified global states occur due to the interplay of spreading through 1D and 3D. (ii) Coexistence between 〈*S*〉 ≈ ±1 requires either a first-order collapse transition [19] or partial second-order collapse [19] of the chromatin. In contrast, in the case *∊*_MM_ = *∊*_UU_, chromatin does not collapse but stable domains form.(iii) The lattice [19] and the off-lattice models [18] explore only the slow-spreading regime. By exploring systematically both the fast- and slow-spreading regime, our model examines the interplay between the time scale of writing (marking) and erasing (unmarking) kinetics and the time scale associated with polymer relaxation time. (iv) One of our most important findings is the importance of the nucleation site in controlling the formation of a stable heterochromatin domain. Finite domains are also found in models (interactions between modified nucleosomes are attractive), which have a number of genomic bookmarks (GBMs) that in effect are the boundary elements, which stop epigenetic spreading [40]. In this model, it is found that finite size domains, with spacing that is given by the genomic distance between the GBMs, form only if the linear density of GBMs exceeds a critical value. The nucleation site was considered in a model [26] by exploring the competition of a large number of densely placed nucleation sites, that participate in spreading two different epigenetic marks.

In contrast, we investigated the effect of spreading a single mark over a stretch of chromatin, without the competition from other competing processes. The combined findings in these and other one-dimensional models, which do not explicitly take the polymer dynamics into account, show that there are several scenarios in the formation of stable epigenetic domains. It is only by quantitatively analyzing specific experiments that we can assess the validity of various models.

### Estimate of *k*^+^

It is instructive to calculate *k*^+^ for the chromatin model to assess the potential relevance of our results to experiments. Because the value of forward rate (*k*^+^) is expressed through the relaxation time of chromatin fiber, *τ_r_*, a realistic estimate can be made using polymer theory. According to the Zimm theory, 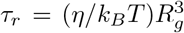 where *η* is the viscosity of the nucleoplasm, *k_B_* is the Boltzmann constant, and *R_g_* is the radius of gyration of the 60,000 bps chromatin fiber investigated in this work. In a good solvent, *R_g_* = *Al_k_N*^0.6^ where *l_k_* is the Kuhn length of a monomer, and *A* ≈ 1/6 for polymer [41]. The persistence length of chromatin fiber is about 1 kbps [42], which implies that *l_k_* is 2kbps, and the number of Kuhn monomers is *N* = 60kbps/2kbps = 30. To estimate the value of *b*, we resort to our previous work [32] in which we found that *b* is 70 nm for a 1.2kbps long chromatin segment. Hence, *b* for the 2kbps long segment is on the order of 100 nm. Combining *b* and *N*, we obtain *R_g_* = *αbN*^0.6^ ≈ 100*nm*. The viscosity of the nucleoplasm is experimentally measured to be 10^3^Pa · s [43]. Using these values, we find that *τ_r_* ≈ 240s. Due to the likely over- or under-estimation in the above calculations, we take the *τ_r_* to be in the range from 10^2^s to 10^3^s. Thus, the forward rate for the slow-spreading in our work is in the range from *k*^+^ = 0.36hr^−1^ to *k*^+^ = 0.036hr^−1^, and the forward rate for the fast-spreading in this work is in the range from *k*^+^ = 3600hr^−1^ to *k*^+^ = 360hr^−1^.

It is worth noting that for eukaryotic H3K9me3 domains, *k*^+^ has been estimated to be 0.1 hr^−1^ - 0.15 hr^−1^ by using a 1D spreading model (see below) [37] to fit ChIP-seq data [44]. Hence, we reason that in this work, the slow-spreading case is potentially relevant to the cell *in vivo* with the fast-spreading case providing a theoretically interesting limit. It is worth noting that the range of the calculated *k*^+^ were done without using any experimental data.

### Finite domain formation through 1D processes

The formation of finite domains seems unlikely when we only allow 1D spreading. However, using essentially a 1D model (nucleation, propagation, and turnover or noise) Hodges and Crabtree [25] showed that finite domains do form, thus rationalizing experimental data [37]. Given that the value of *k*^+^ using the Zimm time for *τ_r_* is in the ballpark of the experimental data [37], we estimated the time needed for the finite domain to be established. From the results in Figure S18 in the SI, which shows the number of modified nucleosomes as a function of time, we find that if slow-spreading is restricted to occur only through 1D process (blue line in Figure S18(B), finite domain does not form even after 2 × 10^5^*τ_r_*. With *τ_r_* ~ 240s the estimated time is on the order of 10^4^ hours, a value that vastly exceeds typical cell cycle time. By comparison, if spreading occurs by 3D (yellow curve in Figure S18(B)), a finite domain is established after ≈ 0.5 × 10^5^*τ_r_*, which is on the order of 10^3^ hours. Thus, in our model 3D spreading is considerably more efficacious, occurring in a finite number of generations.

### Comparison to experiments on Mouse Embryonic Stem Cells

We explored the biological pertinence of our results by comparing to the H3K9me3 enrichment profiles reported elsewhere (see Fig. 6 in [37]). In this study, the authors investigated the propagation of HP1*α*-induced of H3K9me3 modification in the 10 kb *Oct4* locus in embryonic stem (ES) cells and Mouse Embryonic fibroblasts (MEFs) [37], and found that in both the ES and MEF cells H3K9me3 propagated symmetrically to produce finite spatial domains. To rationalize the observations, a one-dimensional model was proposed [25, 37] in which spreading occurs with finite probability to the neighboring sites from an already modified nucleosome. In addition, turnover (M → U transition) could occur randomly from any marked nucleosome, which clearly opposes spreading. The center nucleosome in the 1D lattice represents the site of csHP1*α* (analogue of the NS in our models), containing the domain that specifically nucleates the H3K9me3 mark. By tuning the ratio 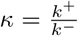 they could produce the small (large) domain sizes in the MEF (ES) cells. In order to rationalize the validity of the 1D model model, they compared their results to ChIP-seq experiments, which reported [44] enrichment in H3K9me3 marks in Mouse Embryonic Stem Cells. The study [44] used chromatin immunoprecipitation followed by sequencing is performed. The results are equivalent to a temporal average of an idealized experiment with spatio-temporal resolution. By means of k-means clustering [37] it was found that the data for ES cells [44] partitioned into two groups; small domains that encompass 77.1% of the data (Figure 8, left panel) and large domains that account for the rest (Figure 8, right panel) [37]. Using the predicted epigenetic profile, they extracted the two parameters (*k*^+^ and *k*^−^) for the small and large domains.

**Figure 8:**
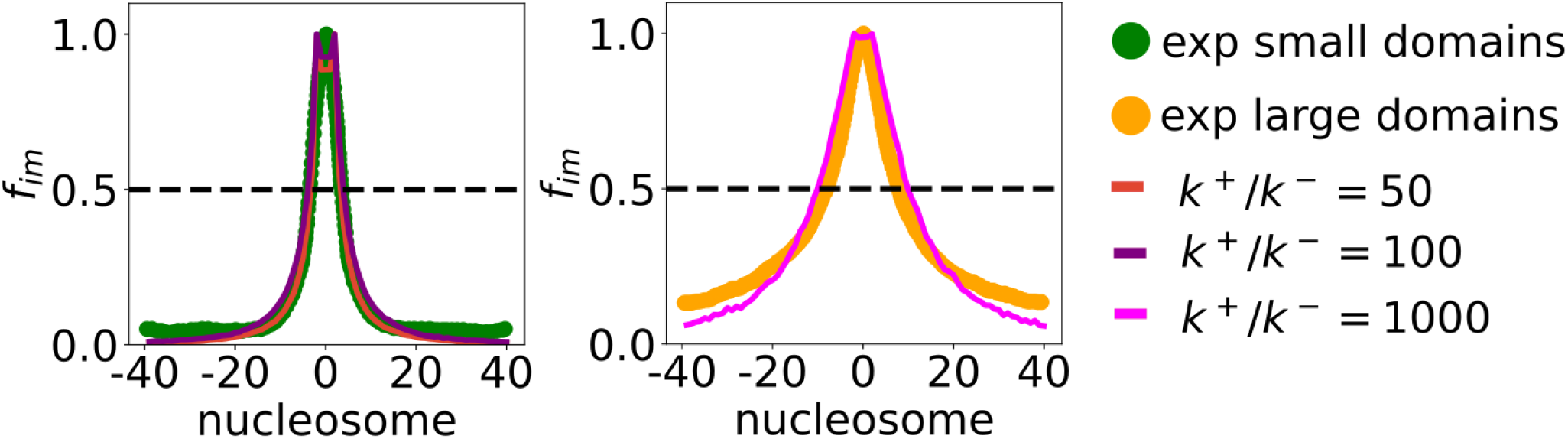
Comparison between simulations performed using Scheme **II** in the slow spreading regime at different *k*^+^/*k*^−^ values with experimental data for H3K9me3 domains in embryonic stem cells [44] that was analyzed using the 1D model [37]. For the small domains, a range of *k*^+^/*k*^−^ reproduces the data (left panel). A higher *k*^+^/*k*^−^ = 10,000 value is needed to account for the larger domain size shown in the right panel. The parameters are chosen to be *k*^+^ = 0.01/*τ_r_* and *α* = 2*k*^−^/*k*^+^.

In order to assess if our model, which involves an interplay of 1D and 3D spreading, could be used to reproduce the reported H3K9me3 enrichment profiles described above, we performed calculations using Scheme **II**. Using *k*^+^ = 0.01/*τ_r_*, we find that for a range of *k*^+^/*k*^−^ values, we obtain good agreement with the enrichment profiles reported using the 1D model (Figure 8), for the both small and large domains. The results in Figure 9 show that other mechanisms like the one we have developed could also explain biologically relevant data for spreading. Of course, this requires different values for the modification rates as well as probabilities for spreading by 3D looping mechanism.

**Figure 9:**
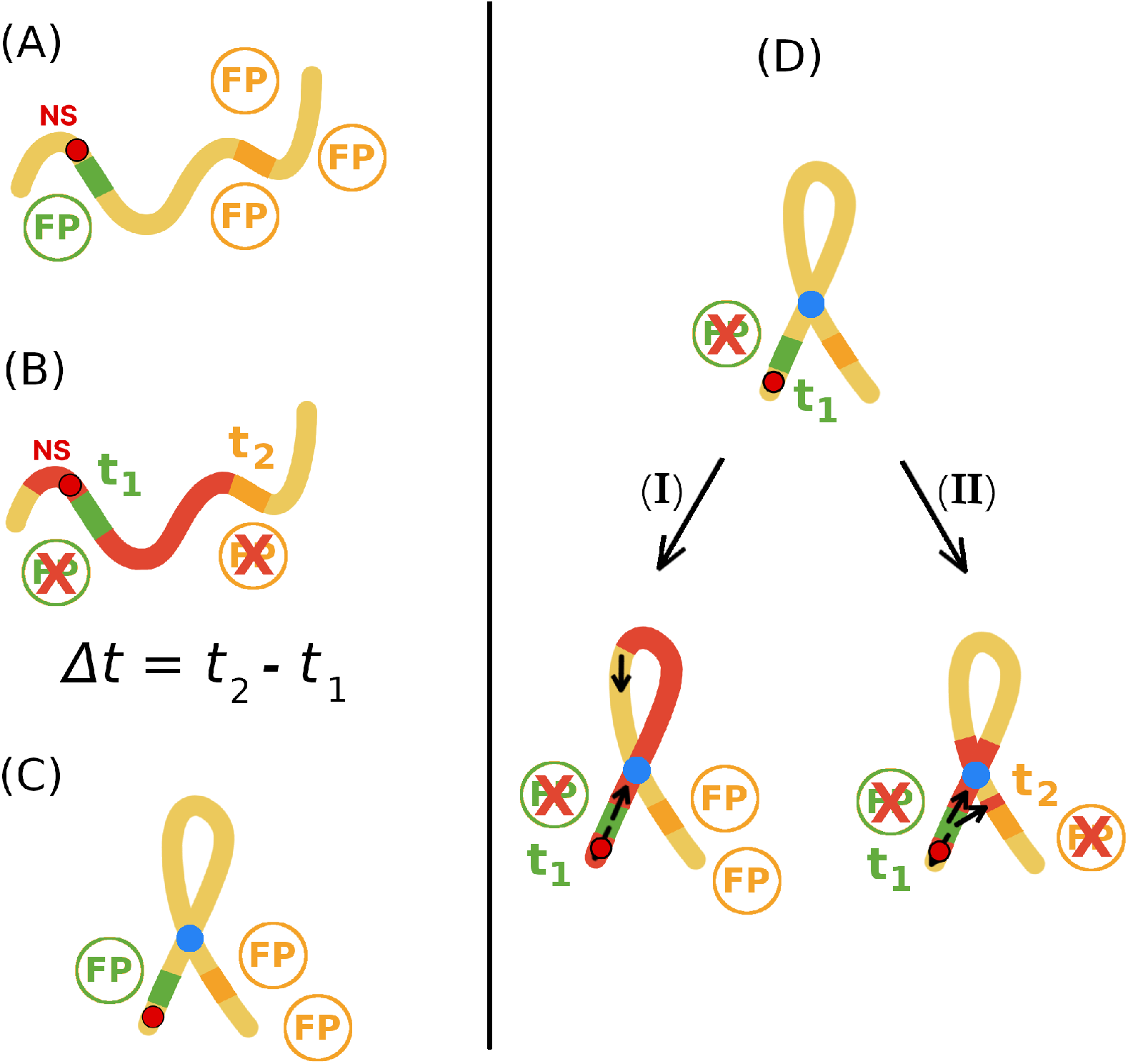
Scheme for the proposed experiment. (A): The gene for the orange fluorescent protein (OFP, in orange) is expressed if chromatin is unmarked (yellow). The repression of the green fluorescent protein gene (GFP) acts as a proxy for the binding of epigenetic spreading initiators to the NS, shown in red circle with black border. Because the nucleation event has not occurred, the GFPs and OFPs are produced. The presence of fluorescent proteins (FP) could be detected by measuring fluorescence intensity at a given wavelength. (B): Binding of epigenetic spreading initiators to the NS represses the GFP expression at time *t*_1_, which could be measured. Subsequently, epigenetic markers (red) spread until the OFP gene is inactivated, resulting in the disappearance of orange fluorescence at *t*_2_, which is measurable. During spreading, chromatin can form loops, but need not have a permanent structure. The time difference, Δ*t* = *t*_2_ - *t*_1_, could be measured for a number of single cells to obtain the distribution, *P*(Δ*t*). This represents the control experiment; similar experimental setup has been employed in Ref [47]. (C): Permanent loop (blue) in chromatin structure may be constructed so that the spatial proximity of the NS and OFP gene is decreased for a given genomic NS-FP distance. (D): Using the FPs as reporters for spreading, only cells that lack green fluorescence are taken into account, as it marks the successful initiation of epigenetic spreading from the NS. If the heterochromatin spreading from the nucleation site is strictly 1D then *P*(Δ*t*) should not differ from the result in the control experiment. If 3D spreading is relevant, *P*(Δ*t*) would differ compared to the control experiment. The same conclusions should be achieved by placing the OFP gene at different genomic distances from the NS, for a given loop length.

The good agreement shown in Figure 9 allows us to draw a few conclusions. (i) In our model as well as in the 1D model [37], the NS is situated in every epigenetic domain. Unlike in our model, the probability of the NS getting modified in the 1D model is less than unity, although it has to around 0.75 to obtain stable large domains (Figure 2 in [25]). (ii) In both the models competition between the forward and the backward rate is crucial to account for epigenetic domains of different widths. For example, if *κ* is large domains without bound form. (iii) Importantly, unlike in the 1D model, in our model spreading rate from the NS is greatly enhanced compared to spreading from modified nucleosomes. Nucleation sites are sequence-defined DNA fragments to which enzymes bind with high specificity and it is possible that different conformational changes might arise by binding to different sequences, thus modulating the enzyme activity. It is also possible that different mechanisms might be operative in heterochromatin formation in different species. The combination of 1D and 3D models, which integrates the rates of spreading and chromatin dynamics created here, could be a step towards fulfilling this goal. Indeed, there is experimental evidence pointing to the importance of both linear (nearest-neighbor) [45] spreading of H3K9me3 marks, as well as spatial spreading [46] of H3K27 marks, showing that chromatin silencing could proceed by both mechanisms, as accounted for in Scheme **II** of our model.

### Experimental prospects

To test the prediction that stable finite domains can be driven by 3D-organization of chromatin without boundary elements, we propose an experiment based on fluorescent probes, similar to the one reported in Ref [47]. The difference would be that the nucleation element and position-dependent fluorescent probe are placed within a permanent artificially engineered loop. The proposed experiment is schematically shown in Figure 9, and is explained in the figure caption. If the experiment is successful, it would help elucidate the extent to which chromatin looping plays a role in the formation of epigenetic domains, and more generally whether epigenetic domain formation occurs by linear or spatial spreading, or a combination of both the mechanism.

## Data Availability

The data can be generated using the in-house computer code that has been deposited in Github. The URL is https://github.com/Katamar/epigenetic.

## Acknowledgements

We are thankful to Bassem Al-Sady and Ilya Finkelstein for advice and useful discussions. We also appreciate the comments from Dan Jost and thank him for pointing out a couple of relevant references. This work was initiated while MK was a postdoctoral fellow at the University of Texas.

