## Supplementary Information for "Chromatin dynamics controls epigenetic domain formation"

January 6, 2021

### 1 Langevin dynamics simulations

We describe the time evolution of the chromatin by the Langevin equation,

$$m_i \frac{d^2\mathbf{r}_i}{dt} = \mathbf{F}_i - \xi \frac{d\mathbf{r}_i}{dt} + \mathbf{R}_i(t), \tag{1}$$

where $\mathbf{r}_i$ is the position of the $i^{th}$ locus whose mass is $m_i$, $\mathbf{F}_i = -\frac{\partial U_T}{\partial \mathbf{r}_i}$ is the systematic force arising from $U_T$ (see the main text), $\xi$ is the friction coefficient, and $\mathbf{R}_i$ is the random force that satisfies the fluctuation dissipation theorem. An in house code was developed to integrate the Langevin equation, with $T = 300K$, using the velocity-Verlet algorithm [1]. After equilibrating the polymer for times that exceed the relaxation time ($\tau_{R_{ee}}$) of the end-to-end vector of the polymer, we performed long simulations ($\gg 10\tau_{R_{ee}}$) so that reliable statistics for computing various quantities of interest could be generated.

### 2 Epigenetic modification algorithm flowchart

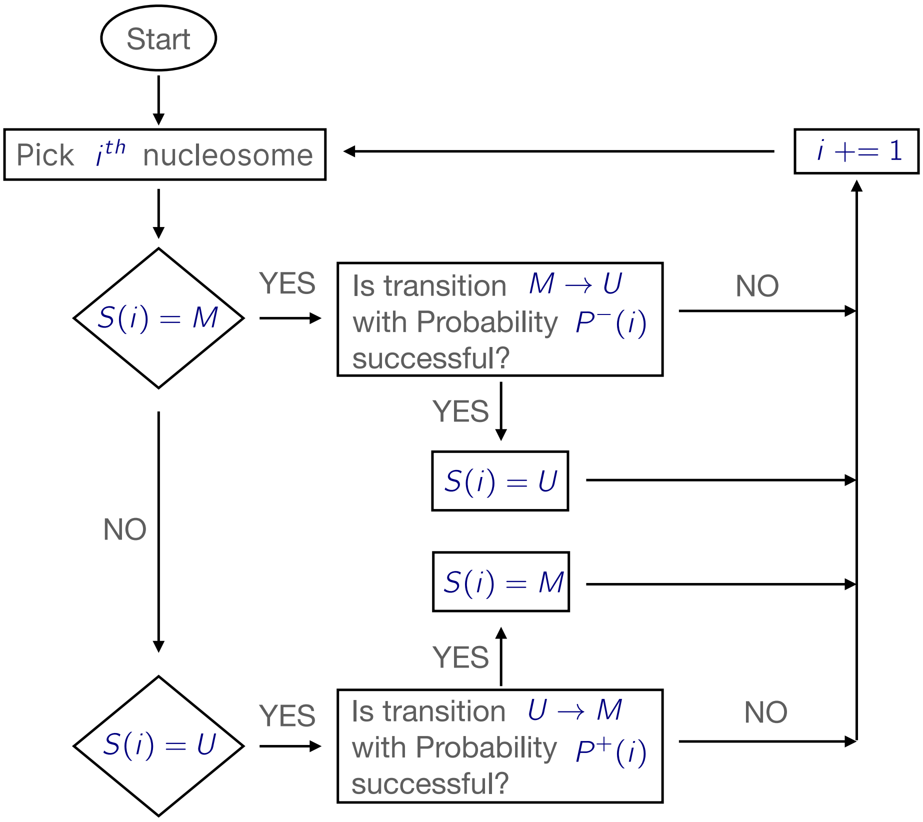


Figure S1: Flowchart of the algorithm for epigenetic modifications at each time step. The probability values are given in Figure 2.

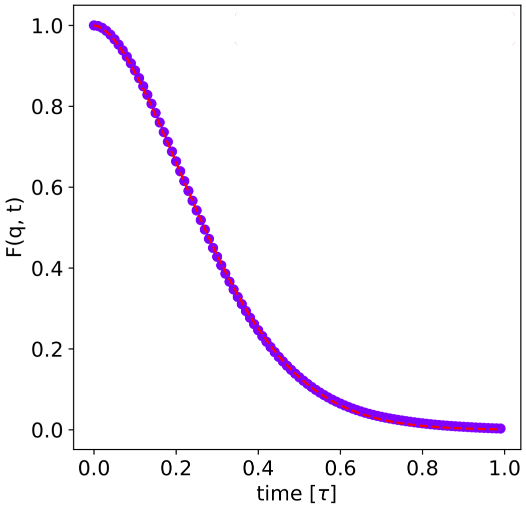


Figure S2: Intermediate scattering function, evaluated at the wave vector $q = \frac{2\pi}{r_c}$ ($r_c = 1.122\sigma$), as a function of time. $F(q,t)$ was averaged over 15 independent trajectories by removing the overall rotational and translational degrees of freedom. The value of $\tau_r$ is extracted using the fit, $F(q,t) = \exp^{-(t/\tau_r)^\beta}$ with $\beta$=1.7, yielding $\tau_r \approx 0.3\tau$ where $\tau$ is the natural time governing Eq. 1.

### 3 Structural relaxation time

We calculated the structural relaxation time, $\tau_r$, from the decay of the structure factor,

$$F(q,t) = \frac{1}{N}\left\langle \sum_{j=1}^{N} \exp^{-iq\cdot(r_j(t)-r_j(0))} \right\rangle, \tag{2}$$

with $q = \frac{2\pi}{r_c}$. In the above equation, $r_j(t) - r_j(0)$ is the displacement of nucleosome $r_j$. From the decay of $F(q,t)$ (Figure S2) we estimated the characteristic time, $\tau_r$. The time $\tau_r$, which is an estimate for the relaxation of the chromatin polymer, is assumed to set the over all time scale. All other rates that are relevant to epigenetic spreading are set relative to $\tau_r$. It is natural to use $\tau_r$, especially for 3D spreading to monitor the modification process. Note that $\tau_r$ is a function of $N$, $l_p$, as well as solvent quality.

### 4 Solvent quality

The dimensions of the flexible chromatin polymer is determined by the Lennard-Jones interaction strength, $\epsilon$ (Eq. 3 in the main text), between the nucleosomes that are separated by at least two bonds from each other. The parameter $\epsilon$ is an effective interaction strength averaged over the environmental factors (solvent, ions, crowding agents etc.) The dimension of the chain (compact or random coil) depends on the second virial coefficient,

$$v_2 = 2\pi \int_0^\infty r^2 dr[1 - \exp^{-\beta U_{LJ}(r)}], \tag{3}$$

where $U_{LJ}(r)$ is given in Eq (3) in the main text, $\beta = 1/k_BT$. If $v_2 > 0 (< 0)$, then the chromatin could be extended (random coil). In Figure S3, we show $v_2$ as a function of $\beta\epsilon$. The $\theta$-point at

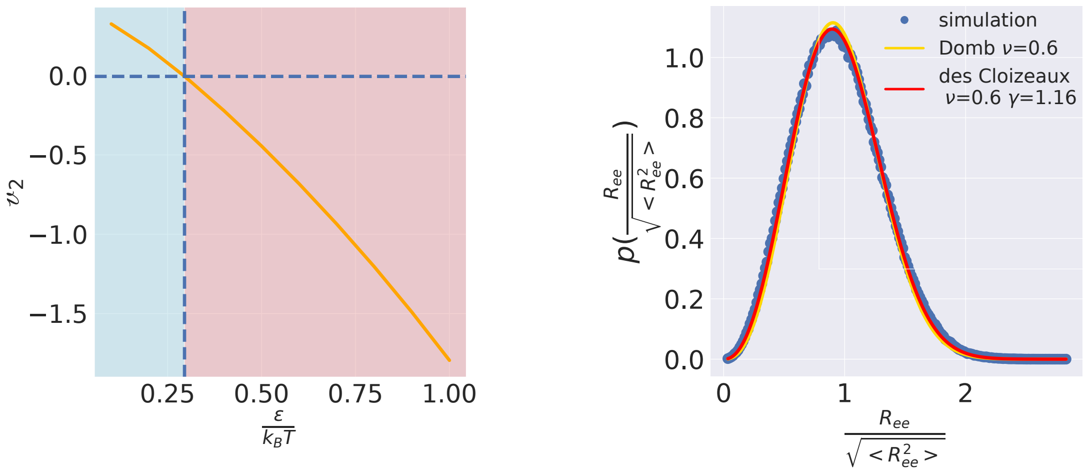


(a) (b)

Figure S3: (a) Second virial coefficient as a function of the Lennard-Jones parameter $\epsilon$, characterizing the interaction between the nucleosomes. The blue (red) region corresponds to "good solvent" ("poor solvent"). (b) Comparison of the end-to-end distance distributions computed from simulations ($\epsilon = 0.1 k_B T$) with rigorous theoretical prediction for a polymer in a good solvent [2].

which $v_2 = 0$ corresponds to $\epsilon = 0.3 k_B T$ (Figure S3). We chose $\epsilon = 0.1 k_B T$, which is in the good solvent region. Thus, the model chromatin behaves as a Flory random coil.

If the chromatin polymer is a random coil, then the distribution $P(x)$, with $x = R_{ee}/ < R_{ee}^2 >^{1/2}$, should follow the universal behavior predicted by polymer theory. In particular, we expect $P(x)$ to be given by $P(x) \sim x^g \exp^{-x^\delta}$, where $\delta \approx 1/(1-\nu)$, where $\nu$ is the Flory exponent ($\approx 0.6$ in 3D). The excellent agreement between theory and simulation confirms that the chromatin polymer is indeed a random coil. We note in passing that the spreading process could change dramatically if $\epsilon$ is changed.

### 5 Contact times and $\tau_r$

There are three time scales that characterize our epigenetic polymer model (i) The rate of forward reaction rate $k^+$(Eq 4 in the main text), (ii) The second is $k^-$, the backward reaction rate. Both $k^+$ and $k^-$ are defined in the main text. (iii) The third is, $\tau_r$, the chromatin relaxation time. Below we describe how these timescales are chosen, and assigned physical meaning.

I As shown in Figure 1, epigenetic spreading could occur either linearly (1D) ($i$ and $i \pm 1$) or by non-bonded nucleosomes that come into proximity through loop formation (3D). The looping time could be substantial, making the modification probability through the 3D mechanism less efficient than by 1D. In order to account for the separation in time scales, we allow for 1D modifications to occur on time scale $t = \gamma \tau_r$. In other words, the probabilities of modification through 1D mechanism are computed using equations in Figure 2 (b). Spreading in 3D occurs if two loci, separated by at least 2 bonds, come into contact. There is a spectrum of looping times that depend on the separation $|i - j|$ between the nucleosomes. A relevant time

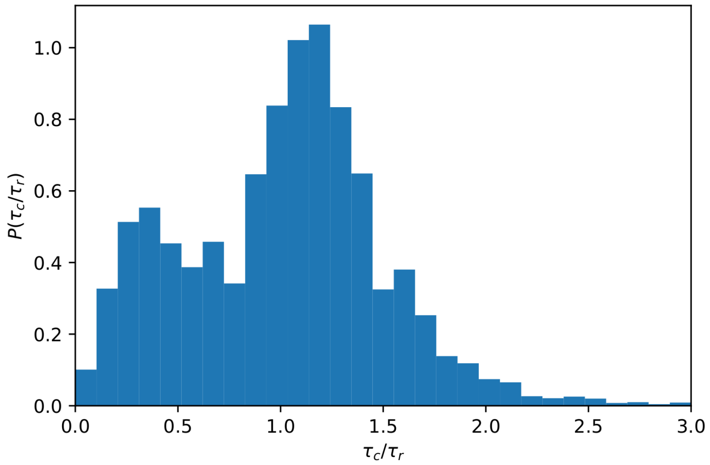


Figure S4: Distribution of contact duration times $\tau_c$. A contact forms if two nucleosomes are within $r_c = 1.122\sigma$ The mean value is $\langle \tau_c \rangle = 0.84\tau_r$.

for modification is the contact life time ($\tau_c$) during which 3D spreading could occur. The distribution of $P(\tau_c/\tau_r)$ contact life times, expressed in units of $\tau_r$, (Figure S4) shows that the average $< \tau_c > \approx 0.84\tau_r$. To simplify the computations, we calculated the probabilities of 3D spreading using equations shown in Figure 2 (b). The use of $\langle \tau_c \rangle$ as a proxy for looping, independent of the genomic separation between the loci, simplifies the computations without qualitatively altering the results.

II We consider two extreme scenarios: fast spreading, which is achieved by using $k^+ = \frac{100}{\tau_r}$. Fast spreading essentially diminishes the 3D mechanism, thus emphasizing 1D processes. Slow spreading occurs when $k^+ = \frac{0.01}{\tau_r}$.

III The ratio $k^+/k^-$ is a free parameter. By covering a range of $k^+/k^-$ values, different scenarios for epigenetic spreading may be anticipated, resulting in chromatin switching from unmodified to modified state, $< S >= -1 \leftrightarrow < S >= +1$. The forward rate, distal from the nucleation site, is $\alpha k^+$ with $\alpha$ less than unity. Different choice of parameters ($\alpha$, $k^+$, $k^-$) produces distinct switching patterns (Figure S7). We chose $k^+/k^- = 50$ in most of the simulations. We also did simulations with $k^+/k^- = 10,000$ in order to assess the effect of drastically accelerating the forward spreading reaction.

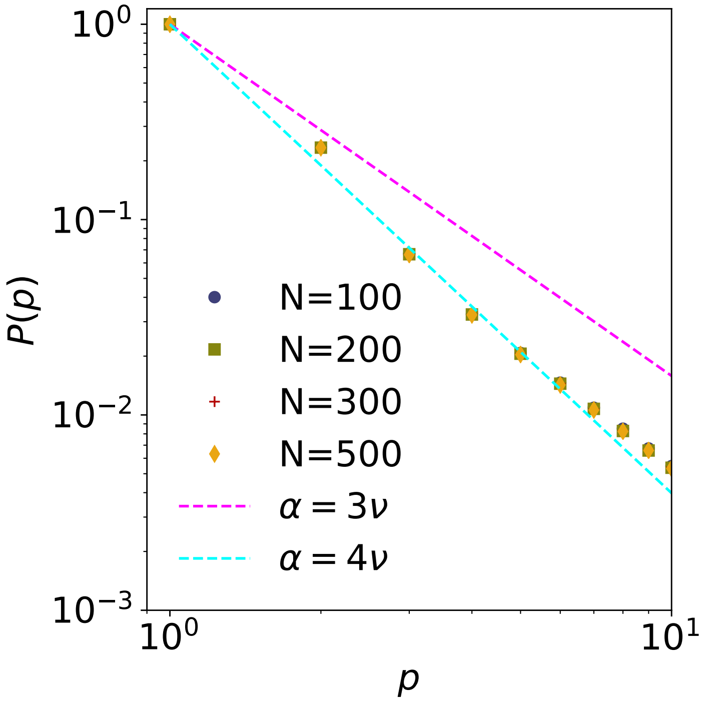


Figure S5: Dependence of the contact probability $P(p)$ as a function of the number of bonds that separate two nucleosomes, $p$. The dashed lines represent theoretical scaling $P(p) \sim p^{-\alpha}$, where $\alpha$ is a multiplicative value of the Flory exponent $\nu = 3/5$.

### 6 Epigenetic ergodicity

In order to asses if the time scales governing spreading result in a non-equilibrium state, we introduce the epigenetic ergodicity measure by generalizing a measure introduced in the context of glasses [3]. Multiple pairs of independent simulations were used to calculate the epigenetic ergodicity, defined as,

$$d(t) = \frac{1}{N}\sum_{j=1}^{N}[s_{a,j}(t) - s_{b,j}(t)]^2, \tag{4}$$

where $s_{a,j}(t)$ is the time-average of the epigenetic state of nucleosome $j$ in trajectory $a$, $s_{b,j}(t)$ is the corresponding quantity in trajectory $b$. If the system is ergodic, each trajectory would explore the entire epigenetic phase space, and the time-average for both the trajectories should converge at long times. Thus, in an ergodic, or quasi ergodic system, the quantity $d(t)$ should vanish at long $t$. Figure S6 shows that $d(t)$ vanishes at long times for all the spreading processes on time scales that are far less than the time needed for steady state spreading to be established. Thus, for the range of time scales considered here, and for $N = 300$, epigenetic ergodicity is established. It is conceivable that epigenetic ergodicity could be broken, resulting in glass-like epigenetic states for different $N$, and solvent quality governing the spreading dynamics. It is unclear if this could confer any biological advantage for epigenetic memory.

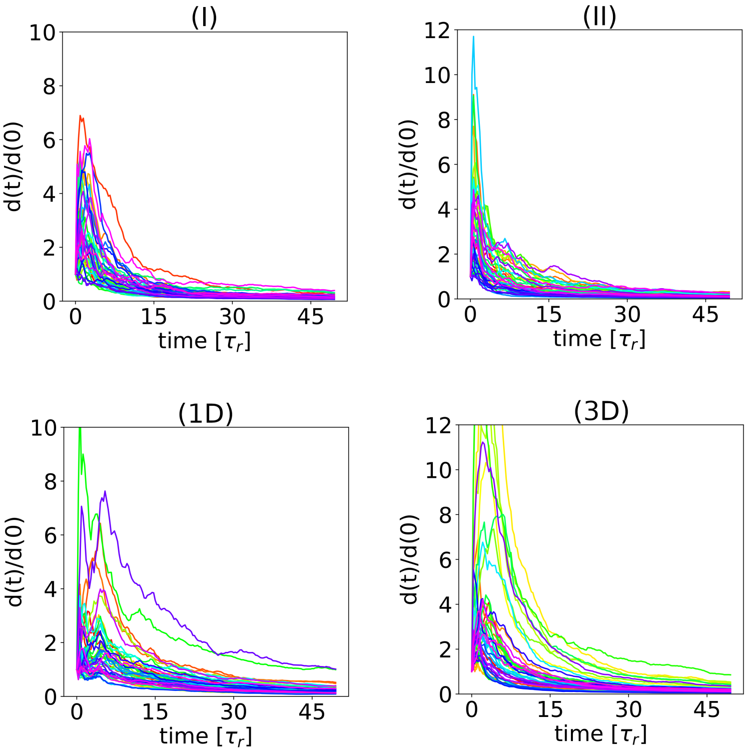


Figure S6: Epigenetic spreading is ergodic. We compared 10 independent trajectories for each biological mechanism. The epigenetic ergodic measure vanishes at long times. The parameter values in simulations are $k^+/k^- = 50$, $k^+ = 100/\tau_r$, with total simulation time $15,000\tau_r$.

### 7 Epigenetic heat map representation

Figure S7 shows a different representation of the data in Figure 2 and Figure S8. The results in Figure S7 show $< S >$ calculated by averaging over 10 trajectories. This data was re-arranged as follows: (i) For a given $k^+/k^-$, $\alpha$ values are rescaled to $\alpha k^+/k^-$. This yields a different set of $\alpha k^+/k^-$ values for each $k^+/k^-$. (ii) With the new scale, $\alpha k^+/k^-$, only $< S >$ data that falls within the range of $\alpha k^+/k^- \in [0.5, 3.0]$ is taken into account. For each $k^+/k^-$ ratio separately, the data is divided into bins with width 0.5. (iii) $< S >$ values are associated to $\alpha k^+/k^-$ bins. If multiple $< S >$ values are in the same, their average is computed. (iv) Each $\alpha k^+/k^-$ bin is associated with a single $< S >$ value, as shown in left panels of Figure 2 and Figure S8.

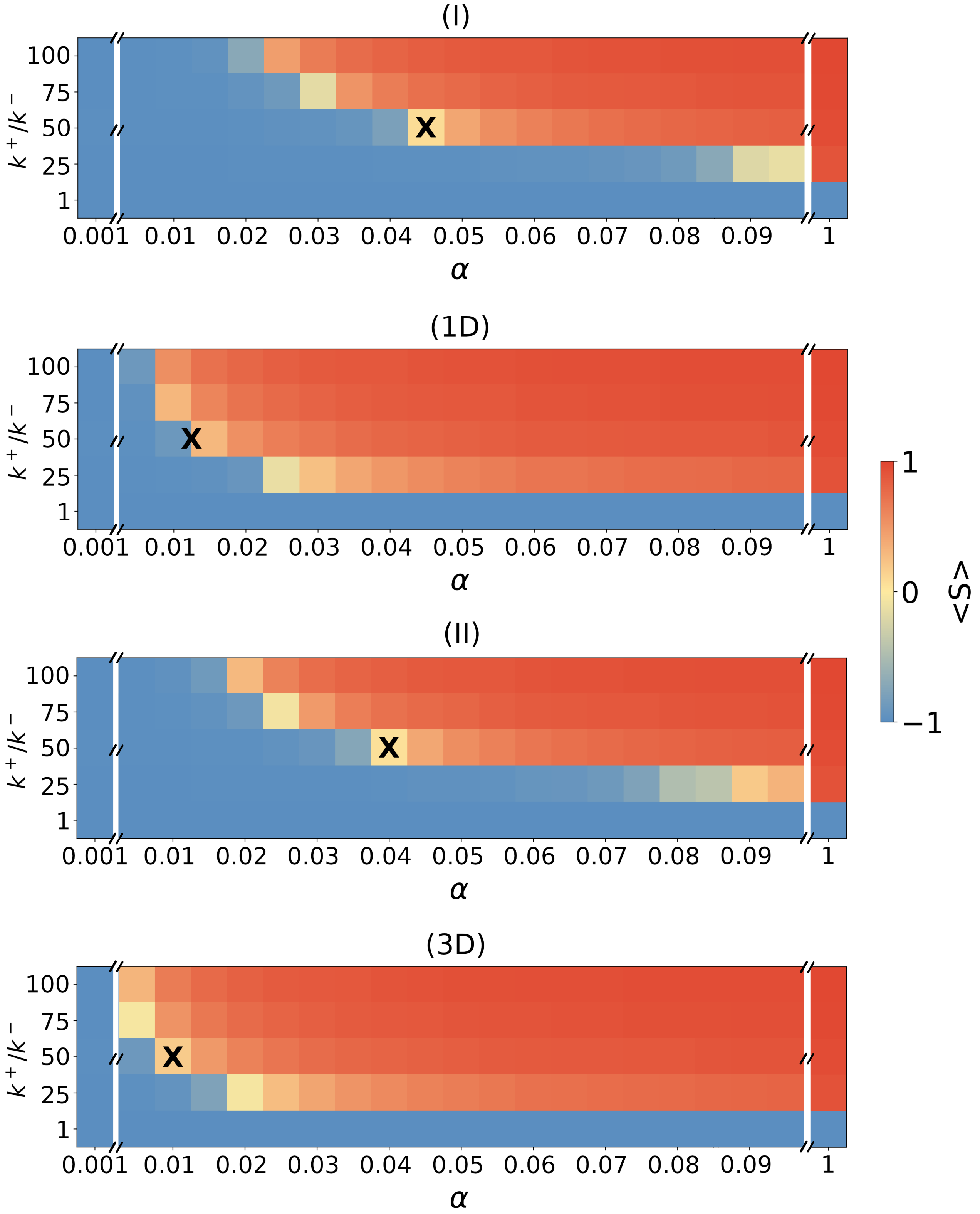


Figure S7: Epigenetic switching shown for all four biological mechanisms as a function of $\alpha$ (Figure 2), quantified using the mean spin value $< S >$. $k^+ = \frac{100}{\tau_r}$, the total length of the simulations was $15,000\tau_r$, while $k^+/k^-$ and $\alpha$ are varied. Note that (1D) and (3D) are limits of (I) and (II), respectively.

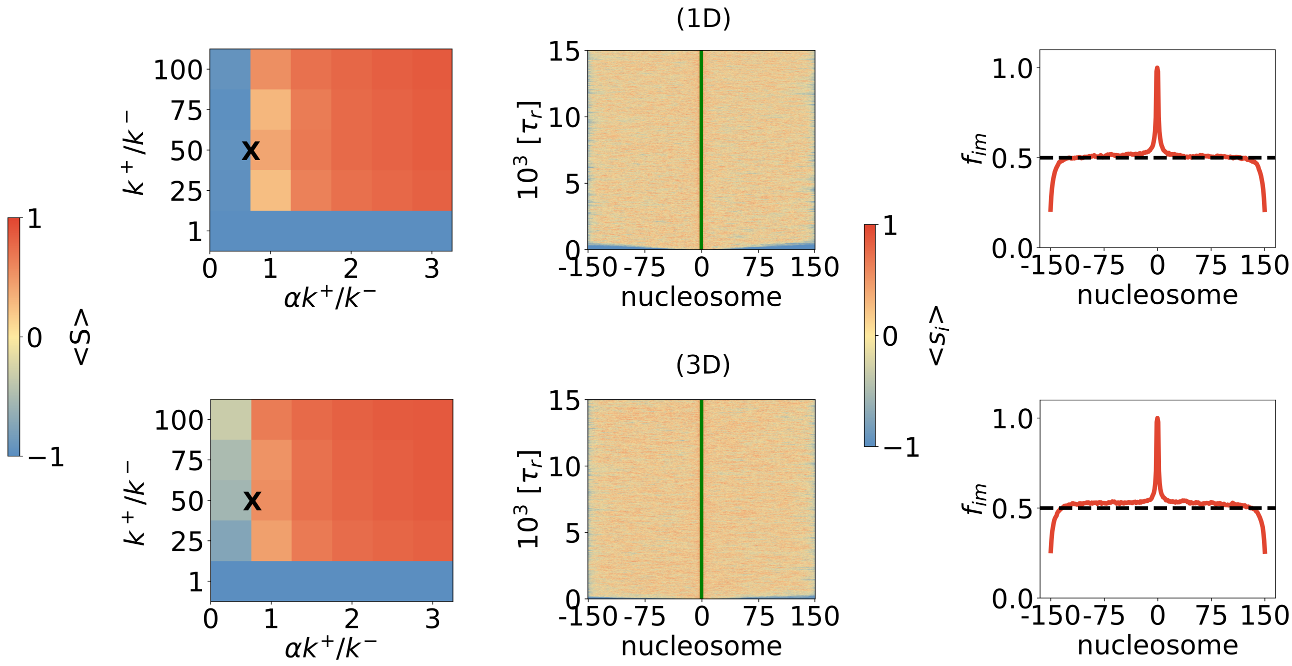


Figure S8: Epigenetic switching depends on the ratio of the forward to backward rate associated with loci that are distal to the nucleation site. Left panels: epigenetic state is determined by the value of the order parameter $< S >$. Middle panels: average epigenetic state of individual locus $< s_i >$, where $< S > \approx 0$, corresponding to $\alpha = 0.63k^-/k^+ = 0.0126$ for 1D and $\alpha = 0.45k^-/k^+ = 0.009$ for 3D. Right panels: fraction of modification for each nucleosome during the course of the trajectory, $f_{im}$. In all the panels, $k^+ = \frac{100}{\tau_r}$, the total length of the simulations was $15,000\tau_r$, and $k^- = \frac{k^+}{50}$. Vertical green line is the position of the nucleation site.

### 8 Influence of chromatin persistence length on epigenetic spreading

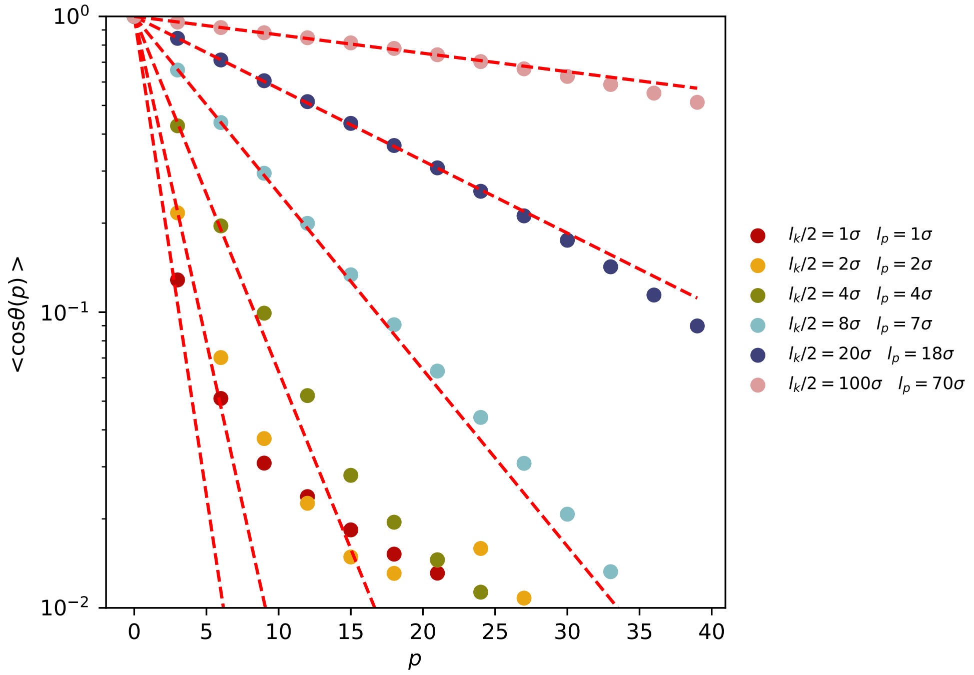


Figure S9: Bond vector correlation function $< cos\theta(p) >$ as a function of the bond distance $p$ along the polymer for different intrinsic stiffness values. Red dashed lines are fits to the curves. The values of the persistence length $l_p$ (listed on the right) are extracted using Eq 5. The fits are increasingly more accurate as $l_p$ increases.

Polymer conformational fluctuations, which affect 3D spreading depends on the intrinsic persistence length, $l_p$. It is difficult to estimate $l_p$ for genomes. It could be argued that $l_p$ may not even be uniform within a single chromosome. Nevertheless, one can anticipate two extreme scenarios. If $L/l_p >> 1$ ($L$ is the contour length of the polymer) then 3D transient loop-driven spreading, as found here and elsewhere (Ref [4], [5]), is likely. In the opposite stiff chain limit $L/l_p << 1$, we expect that spreading would be predominantly determined by the 1D mechanism. These predictions are based solely on the equilibrium polymer characteristics of chromatin without regard to the modification rates.

In order to verify these expectations, we performed simulations by varying the bending stiffness ($l_k$) in the bond angle potential. For different values of $l_k$, we extracted the persistence length using,

$$\langle cos\theta(p)\rangle = \exp^{-\frac{p}{l_p}}, \quad (5)$$

where $\theta$ is the angle between two bond vectors separated by a distance $p$ along the contour of the polymer. The exponential fit to simulations, with $l_k$ ranging from $(2 - 200)\sigma$, is shown in Figure S9. A crossover between the flexible chain ($L \gg l_p$) to a rigid chain is expected when $L \ll l_p$. Figure S9 shows that the best fit is obtained when $l_p$ spans several bond distances, while

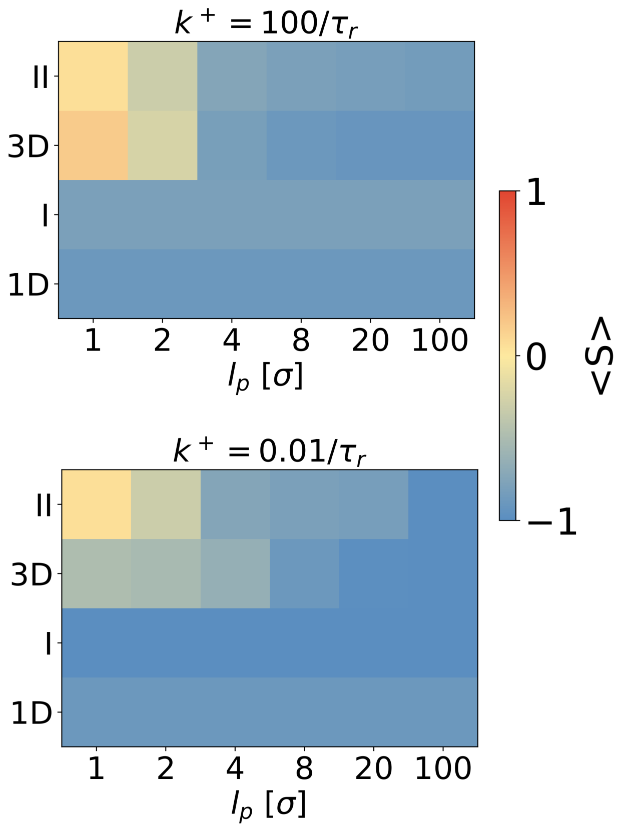


Figure S10: Heat maps showing the average value of the epigenetic order parameter $< S >= \frac{n_m - n_u}{N}$ as a function of the persistence length. It is clear that the 3D result converges to the 1D result as the persistence length increases.

the rigid and flexible limit exhibit stronger deviations from the nominal estimate, $l_p = l_k/2$. The exponential decay $\langle cos\theta(p)\rangle$ (Eq. (5)), is valid strictly for a polymer without excluded volume interactions, and the deviations of the exponential scaling have previously been reported for large $p$ when excluded volume conditions are taken into account [6, 7].

We characterize the global epigenetic state using $< S >$ for two values of $k^+$ as a function of $l_p$. As anticipated, the 3D model converges to the 1D limit (Figure S10) once the chain stiffness increases substantially ($\frac{l_p}{\sigma} \geq 8$). For instance, when spreading is fast ($k^+ = 100/\tau_r$) or slow ($k^+ = 0.01/\tau_r$), the value of $< S >$ for I and II mechanisms are roughly the same at $\frac{l_p}{\sigma} = 8$, as shown in Figure S10. Thus, regardless of the enzyme rates for modifying a nucleosome, the 1D and 3D mechansims converge when $l_p$ is sufficiently large. The reason is that in the stiff chain limit, the energy penalty to bend the polymer is high, thus effectively preventing the formation of looping contacts, which is needed for 3D spreading.

Upon closer observation of 3D spreading, the simulated domains in Figure S11 reveal that enhanced flexibility improves the propensity for stable domain formation. In particular, we observe a stabilizing effect of enhanced flexibility on the epigenetic pattern around the nucleation site. These results show that there is an interplay between 3D and 1D spreading, which is determined by $L/l_p$ and solvent conditions.

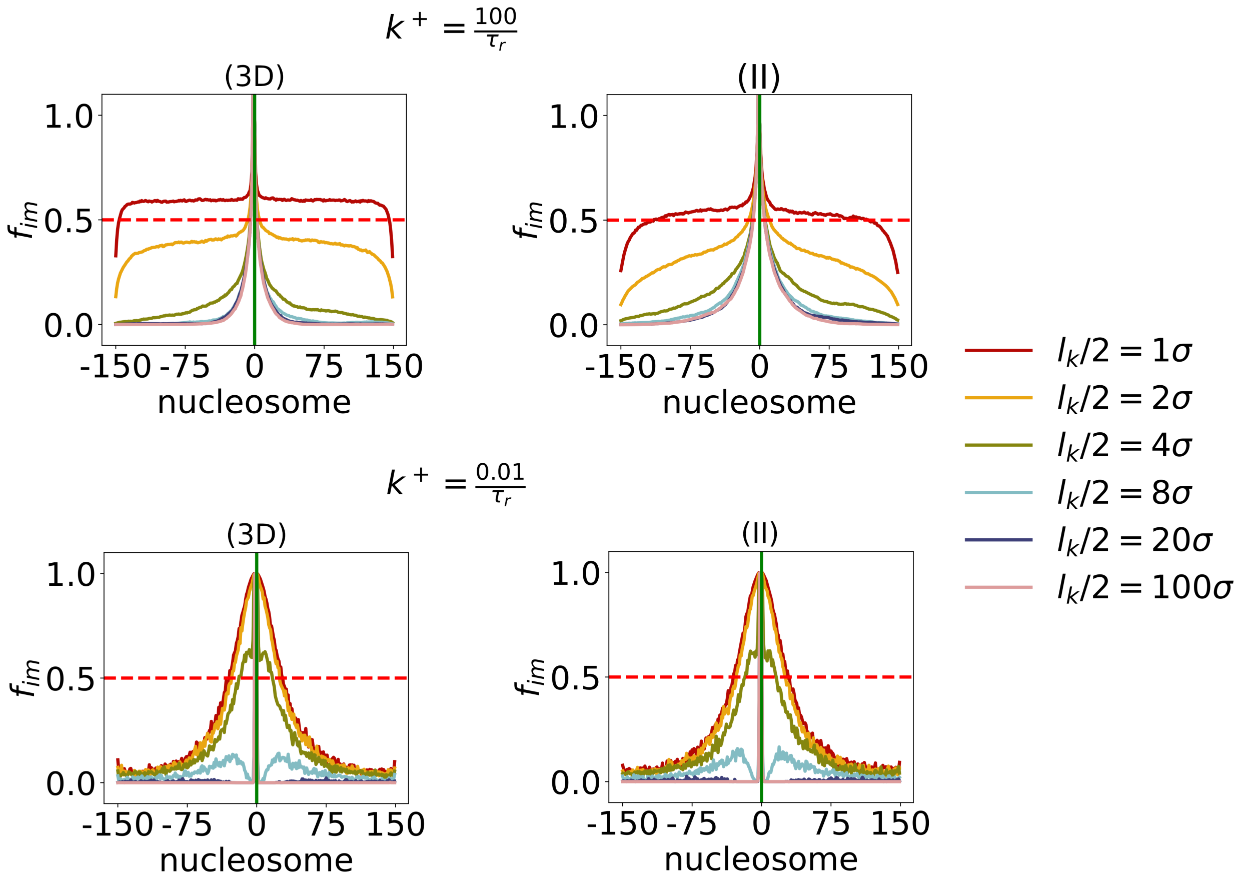


Figure S11: The effect of persistence length, $l_p = l_k/2$, on 3D spreading depicted as the fraction $f_{im}$ of modified state of nucleosome $i$. 3D models are tested under fast, $k^+ = \frac{100}{\tau_r}$, and slow, $k^+ = \frac{0.01}{\tau_r}$, spreading conditions. We used $k^+/k^- = 50$ for fast spreading, and $k^+/k^- = 10,000$, which are the same values used in the main text.

### 9 Finite size effects

Polymer length $N$ affects relaxation timescales, which in turn changes polymer looping kinetics [8], with possible effects on the 3D spreading mechanism. We assessed the effects of changing $N$ from (100-1000) for the fast spreading mechanism. Figure S12 shows that spreading is uniform and exhibits little variations. The main feature in all curves is similar, namely the $f_{im}$ profiles fluctuate around $f_{im} = 0.5$, which is not surprising because the choice of parameters is such that expected value of $< S > \approx 0$. The exception to this are the residues near the nucleation site, which are modified more frequently, and the residues at chain ends, which are modified less frequently. At chain end, the spreading in 1D is not bi-directional, decreasing effectively the probability of spreading. The finite size effect is independent of the spreading mechanism or chain length and affects up to 40 nucleosomes (Figure S13).

In the slow-spreading regime, Fig S14, the 1D spreading shows very little local spreading due to low spreading probability at each time step. However, Fig S14 (II) and (3D) reveal that the inactivation domain profiles behave identically at equal distances from the nucleation site, independent of the length of the polymer. This is because the nucleation site is a major contributor

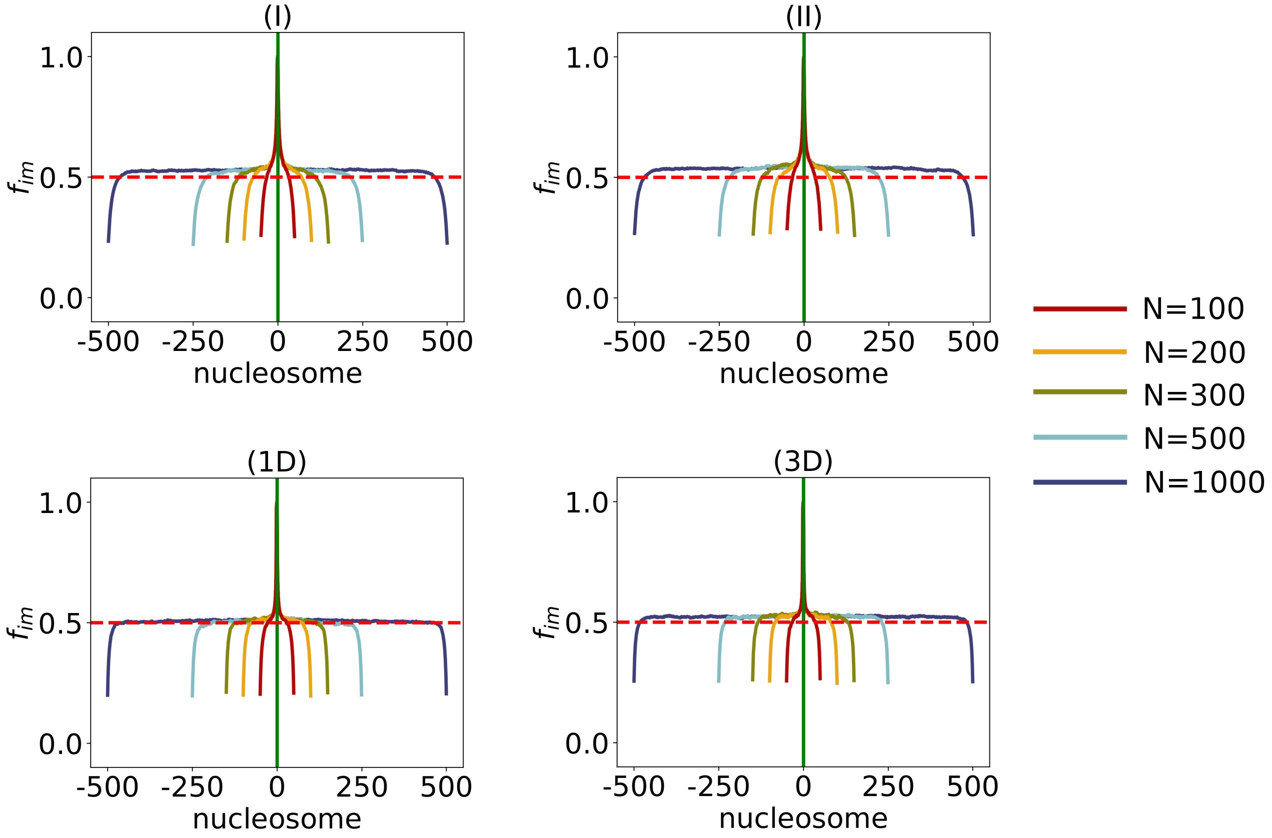


Figure S12: Biological models are tested by varying the locus length, $N$, for the fast-spreading case, $k^+ = \frac{100}{\tau_r}$, $k^+/k^- = 50$.

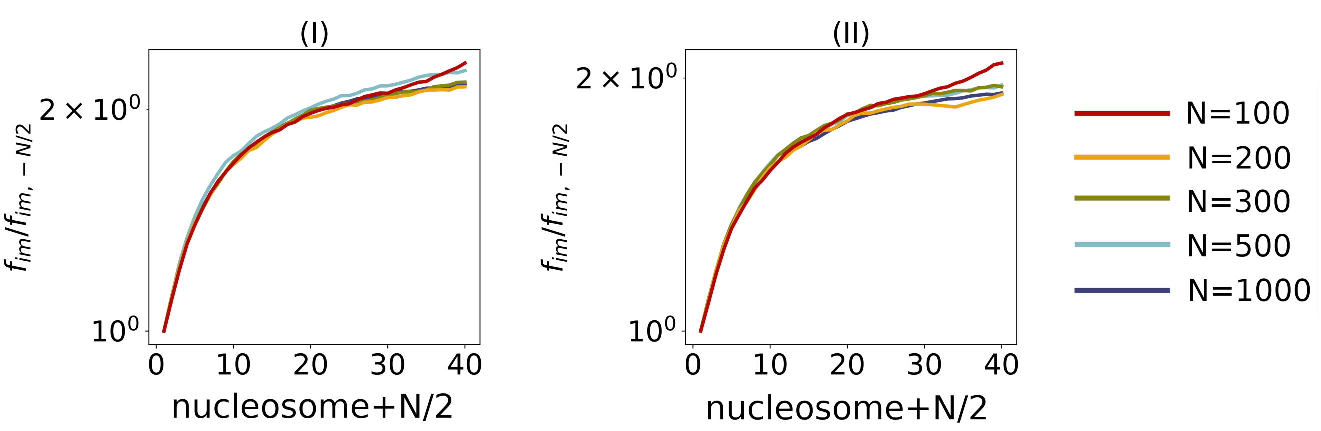


Figure S13: Finite-size effect in the fast spreading regime; region close to end of the chain shows similar patterns of modification for all chain lengths and mechanims I and II. $k^+ = \frac{100}{\tau_r}$, $k^+/k^- = 50$.

to domain formation, and 3D contact formation of nucletion site with other residues determines the domain shape. The fraction of modification decreases with genomic distance from the nucleation site, underlined by lower probability of contact formation at larger distances (Figure S5). Thus, similar domain shapes for chains of different lengths simply reflect a given probability contact scaling with genomic distance, which is constant in these simulations.

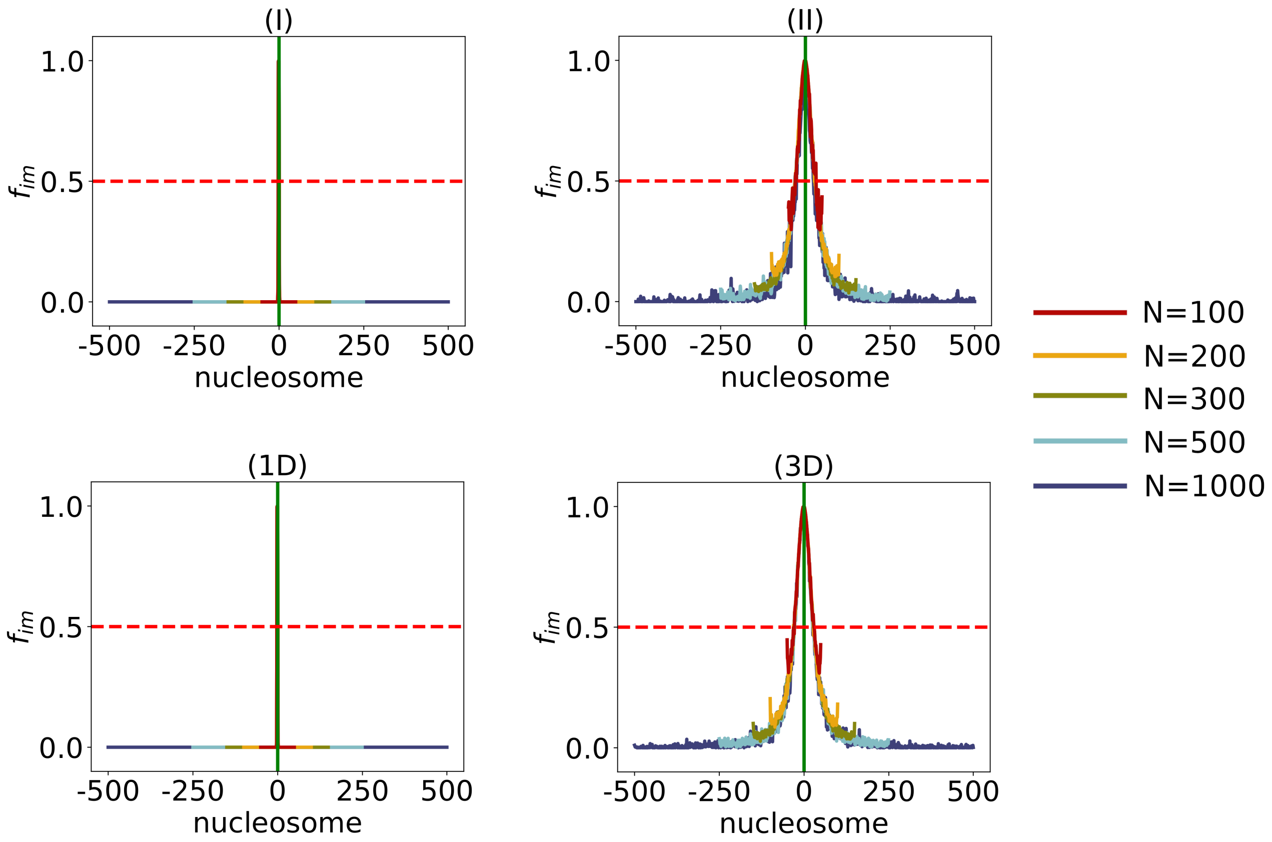


Figure S14: Tests for biological model as a function of locus length, $N$, for the slow-spreading case, $k^+ = \frac{0.01}{\tau_r}$, $k^+/k^- = 10,000$.

### 10 Role of the nucleation site (NS)

At least within the parameter ranges explored in our study, the NS plays a key role in the spreading of the modification. The absence of the NS completely abolishes the possibility of establishing a modified domain, independent of the underlying mechanism (Figure S15). The results were obtained using $k^+/k^- = 50$ and $k^+ = \frac{100}{\tau_r}$. The total simulation time was $15,000\tau_r$.

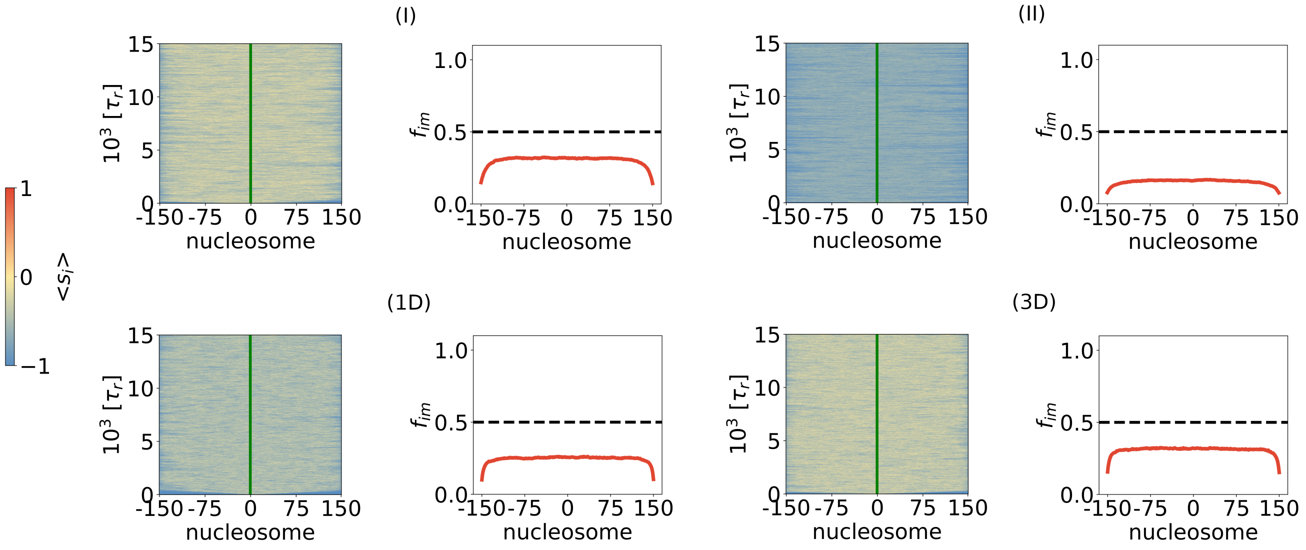


Figure S15: Nucleation site is necessary for domain establishment. Subfigures correspond to four distinct biological mechanisms for spreading. In each subfigure, the left panel is the average state of each nucleosome. The right panel is the fraction of time a nucleosome is in the modified state. The horizontal dotted line represents 50% fraction modification.

### 11 NS location

We investigated the effect of changing the location of the NS on the spreading process. Given that mechanistically spreading occurs bidirectionally from the NS along the chromatin chain, we expected similar results if the NS location is changed. The simulations confirm that it is the NS that drives spreading both in schemes **I** and **II**. The results are similar to the ones in the main text except that the spreading profiles are centered around the NS (Figure S16 and Figure S17).

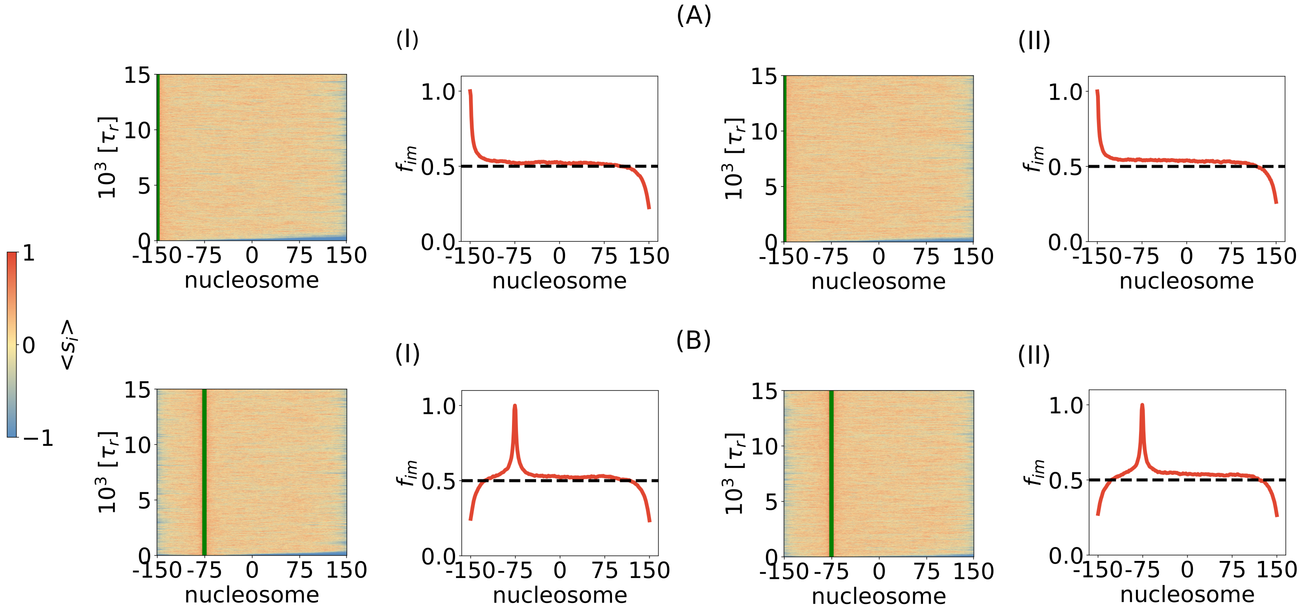


Figure S16: Nucleation site position $i = -150$ (A) and $i = -75$ (B). Both mechanisms I and II are explored. The simulations are performed in the fast-spreading regime.

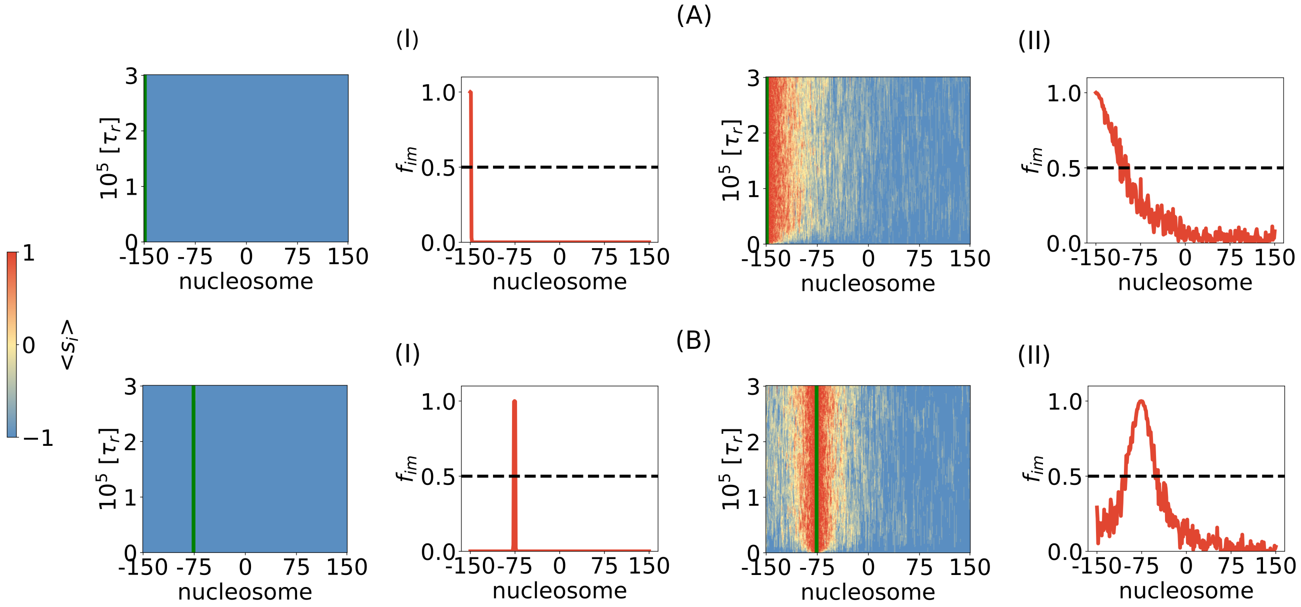


Figure S17: Same as Figure S16 except the results are the slow spreading limit. In (A) , the NS is in position $i = -150$, while in (B) it is in position $i = -75$.

### 12 Time dependence of modifications:

The time needed for modification to be established at each nucleosome site is calculated using,

$$f_{im} = \frac{1}{T}\frac{1}{N_{traj}}\sum_j\sum_t \delta^j_{s_i,+1}, \tag{6}$$

where $\delta^j_{s_i,+1}$ counts the number of occurrences of nucleosome $i$ in state $M$ in trajectory $j$. The time for reaching the global spin state may be obtained using,

$$f_m(t) = \frac{1}{N}\frac{1}{N_{traj}}\sum_i\sum_j \delta^j_{s_i,+1}. \tag{7}$$

The fluctuations in the modified state is given by,

$$\sigma^2(t) = [f_m(t) - f_{av}]^2, \tag{8}$$

where $f_{av}$ is the average of $f_m(t)$ over the $N_{traj}$ trajectories.

There are a few of points that are worth emphasizing using the results in Figure S18. (1) The time needed to reach steady state values in $f_m(t)$ is nearly two orders of magnitude greater in the slow spreading limit than in fast (Figures S18 (A) and (B)). (2) Comparison of $f_m(t)$ in Figures S18 (A) and (B) shows that, at all times, nucleosome modifications occur almost exclusively by mechanism **I** (**II**) in the fast (slow) spreading limits. In the $k^+ = \frac{0.01}{\tau_r}$, there is no possibility of spreading through mechanism **I**.(3) Fluctuations (Eq.8) decay on a much longer time scale in the slow spreading regime relative to fast spreading case (compare Figures S18 (C) and (D)). This is due the importance of modifications occurring exclusively by the looping mechanism. **II** (Figure S18(D)).

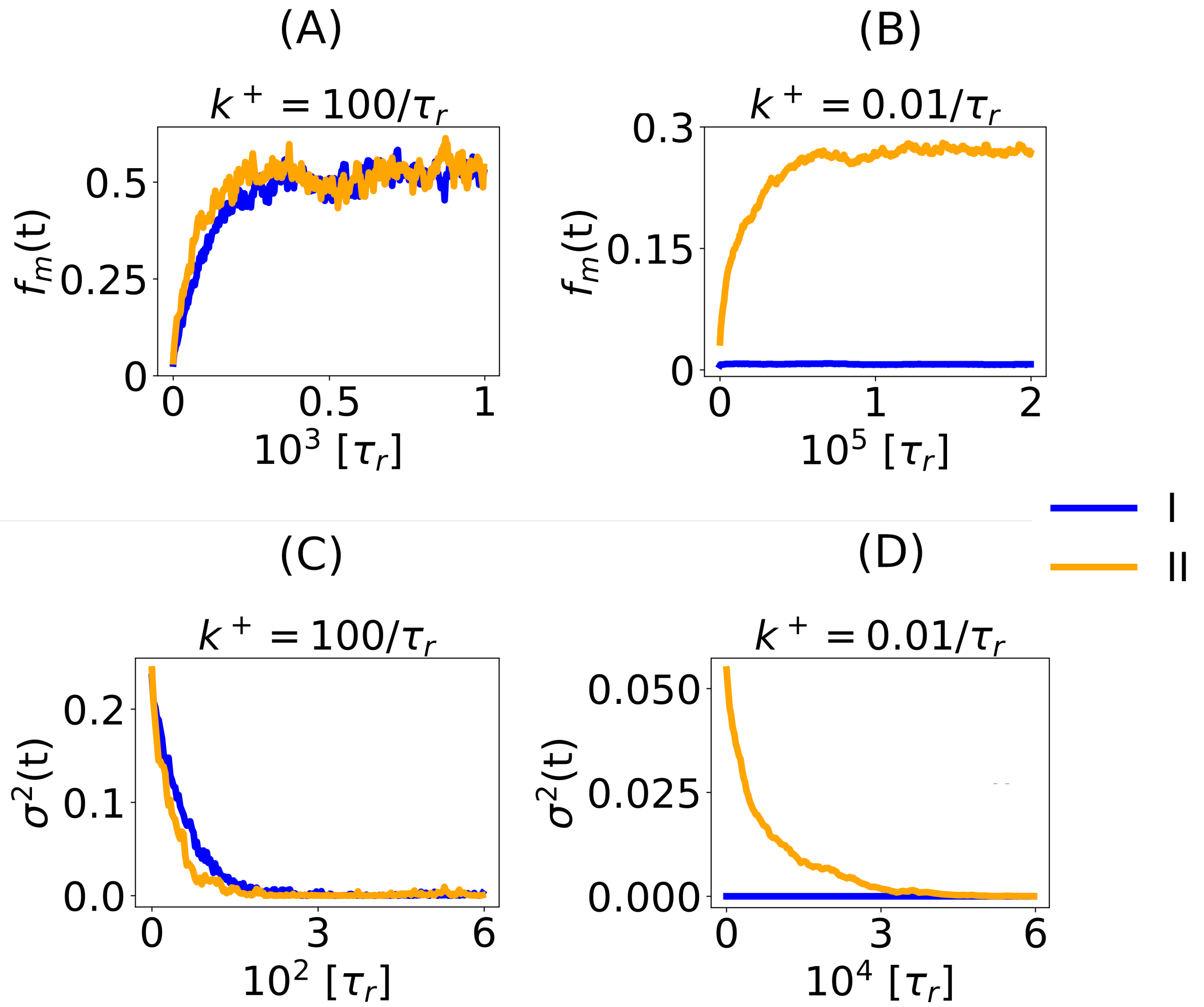


Figure S18: Number of modified nucleosomes as a function of time for mechanisms I and II. (A) Fast spreading (B) slow spreading. (C) and (D) show the results for fluctuations calculated using Eq.8.

[7] Hsiao-Ping Hsu, Wolfgang Paul, and Kurt Binder. Polymer chain stiffness versus excluded volume: A monte carlo study of the crossover towards the wormlike chain model. *Euro Phys. Lett.*, page 28003, 2010.

[8] Ngo Minh Toan, Greg Morrison, Changbong Hyeon, and D Thirumalai. Kinetics of loop formation in polymer chains. *J. Phys. Chem. B*, 112(19):6094–6106, May 2008.
